# Synthetic STARR-seq reveals how DNA shape and sequence modulate transcriptional output and noise

**DOI:** 10.1101/325910

**Authors:** Stefanie Schöne, Melissa Bothe, Edda Einfeldt, Marina Borschiwer, Philipp Benner, Martin Vingron, Morgane Thomas-Chollier, Sebastiaan H. Meijsing

## Abstract

The binding of transcription factors to short recognition sequences plays a pivotal role in controlling the expression of genes. The sequence and shape characteristics of binding sites influence DNA binding specificity and have also been implicated in modulating the activity of transcription factors downstream of binding. To quantitatively assess the transcriptional activity of dozens of thousands of designed synthetic sites in parallel, we developed a synthetic version of STARR-seq (synSTARR-seq). We used the approach to systematically analyze how variations in the recognition sequence of the glucocorticoid receptor (GR) affect transcriptional regulation. Our approach resulted in the identification of a novel highly active functional GR binding sequence and revealed that sequence variation both within and flanking GR’s core binding site can modulate GR activity without apparent changes in DNA binding affinity. Notably, we found that the sequence composition of variants with similar activity profiles was highly diverse. In contrast, groups of variants with similar activity profiles showed distinct DNA shape characteristics indicating that DNA shape may be a better predictor of activity than DNA sequence. Finally, using single cell experiments with individual enhancer variants, we obtained clues indicating that the architecture of the response element can independently tune expression mean and cell-to cell variability in gene expression (noise). Together, our studies establish synSTARR as a powerful method to systematically study how DNA sequence and shape modulate transcriptional output and noise.

## Introduction

The interplay between transcription factors (TFs) and genomically encoded *cis-*regulatory elements plays a key role in specifying where and when genes are expressed. In addition, the architecture of *cis*-regulatory elements influences the expression level of individual genes. For example, transcriptional output can be tuned by varying the number of TF binding sites, either for a given TF or for distinct TFs, present at an enhancer [1, 2]. Moreover, differences in its DNA-binding sites can modulate the magnitude of transcriptional activation, as exemplified by the glucocorticoid receptor (GR), a hormone-activated TF [3–5]. The sequence differences can reside within the 15 base pair (bp) core GR binding sequence (GBS) consisting of two imperfect 6 bp palindromic half-sites separated by a 3 bp spacer. Moreover, sequences directly flanking the core also modulate GR activity [3]. However, these sequence-induced changes in activity cannot be explained by affinity [3]. Instead, the flanking nucleotides induce structural changes in both DNA and the DNA binding domain of GR, arguing for their role in tuning GR activity [3].

Notably, the expression level of a gene is typically measured for populations of cells and thus masks that expression levels can vary considerably between individual cells of an isogenic population [6–9]. This variability in the expression level of a gene, called expression noise, results in phenotypic diversity, which can play a role in organismal responses to environmental changes so called bet-hedging) and in cell fate decisions during development. Expression noise can be explained by the stochastic nature of the individual steps that decode the information encoded in the genome. For example, transcription occurs in bursts [7, 10–12], which can induce variability in gene expression due to differences in burst frequency and in the number of transcripts generated per burst (burst size) [13]. Noise levels are gene-specific, which can be explained in part by differences in the sequence composition of c/s-regulatory elements [11, 14–16]. For instance, the sequence composition of promoters influences expression variability with high burst size and noise for promoters containing a TATA box [15, 17]. In addition, chromatin and the presence or absence of nucleosome-disfavoring sequences have been linked to transcriptional noise [16–19]. Finally, noise levels can also be tuned by the number and by the affinity of TF binding sites [11, 16].

Many fundamental insights regarding the role of sequence in tuning transcriptional output and noise have come from reporter studies [20, 21]. A key advantage of reporters is that they can provide quantitative information in a controlled setting where everything is kept identical except for the sequence of the region of interest. Until recently, a limitation of reporter studies was that sequence variants had to be tested one at a time. However, the recent development of several parallelized reporter assays allows the simultaneous assessment of many sequence variants [21]. One of these parallelized methods is STARR-seq (Self-Transcribing Active Regulatory Region sequencing) [22]. In this assay, candidate sequences are placed downstream of a minimal promoter, such that active enhancers drive their own expression and high-throughput sequencing reveals both the sequence identity and quantitative information regarding the activity of each sequence variant. The STARR-seq method has been used to assay enhancer activity genome-wide [22, 23], to study regions of interest isolated either by Chromatin Immunoprecipitation (ChIP) or a capture-based approach [24, 25], and to study the effect of hormones on enhancer activity [25, 26].

Here, we adapted the STARR-seq method to systematically study how sequence variation both within the 15 bp GBS and in the region directly flanking it modulate GR activity. Specifically, we generated STARR-seq libraries using designed synthetic oligos (synSTARR-seq) with randomized nucleotides flanking the core GBS to show that the flanks modulate transcriptional output by almost an order of magnitude. When grouping sequences based on their ability to either enhance or blunt GBS activity, we found that each group contained a broad spectrum of highly diverse sequences, but striking similarities in their DNA shape characteristics. Using the same approach, we also assayed the effect of sequence variation within the core GBS. Finally, using single cell experiments with individual enhancer variants, we study how the sequence composition of the response element influences expression mean and noise. Together, our studies establish synSTARR-seq as a powerful method to study how DNA sequence and shape modulate transcriptional output and noise.

## Results

### Measuring the activity of thousands of GR binding sequence variants in parallel using the synSTARR-seq approach

To test if we could use the STARR-seq reporter [22] to study how sequence variation of the GR binding site influences GR activity, we first tested if a single GBS is sufficient to facilitate GR-dependent transcriptional activation of the reporter. Therefore, we constructed STARR reporters containing either a single GBS as candidate enhancer, a randomized sequence or as positive control a larger GBS-containing sequence derived from a GR-bound region close to the GR target gene *FKBP5* (Fig. 1A). The resulting reporters were transfected into U2OS cells stably expressing GR (U2OS-GR) [27] and their response to treatment with dexamethasone (dex), a synthetic glucocorticoid hormone, was measured. As expected, no marked hormone-dependent induction was observed for the reporter with the randomized sequence. This was true both at the level of RNA (Fig. 1B) and at the level of the GFP reporter protein (Fig. S1A). In contrast, we observed a robust hormone-dependent activation both at the level of RNA and GFP protein for reporters with either a single GBS or with the larger genomic *FKBP5* fragment (Fig. 1B, S1A), showing that a single GBS is sufficient for GR-dependent activation of the STARR-seq reporter.

**Figure 1.**
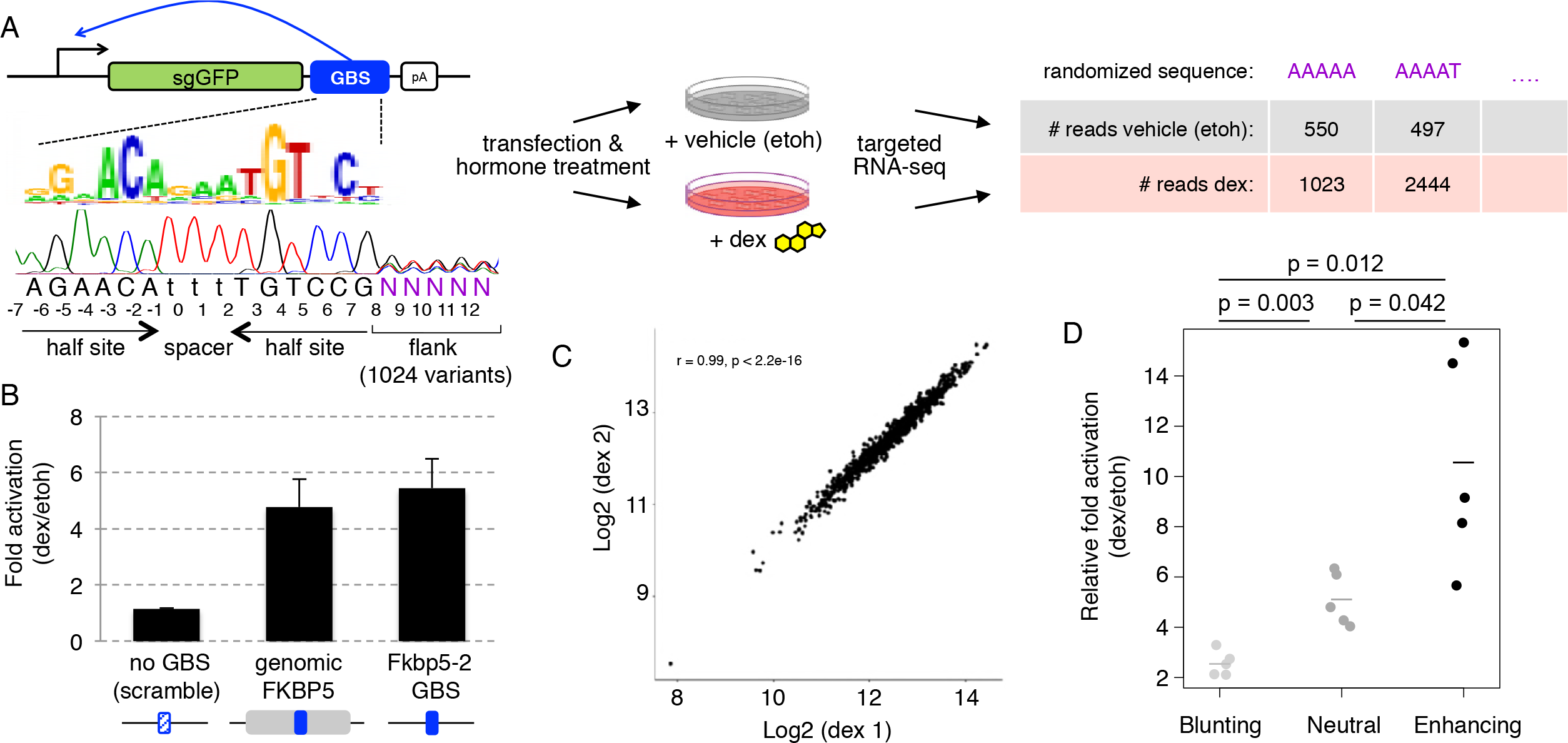
Design and validation of the synSTARR-seq approach. (a) SynSTARR-seq reporter setup using a synthetic library containing a GR Binding Sequence (GBS) flanked by 1024 different flanking sequences (flank library) to screen for flanks that modulate GBS activity. Samples are treated with dexamethasone (dex) or ethanol vehicle (etoh) before targeted RNA-sequencing and counting of the reads. (b) Transcriptional activation of STARR-seq reporter containing candidate enhancer inserts as indicated. Mean fold change upon dexamethasone treatment ± S.D. (n = 3) in U2OS-GR18 cells is shown. Genomic FKBP5 (211bp region hg19Chr6: 35699789-35699999); FKBP5-2 GBS (single GBS: AGAACAtccTGTGCC); no GBS (AGAAACtccGTTGCC). (c) Representative RNA-seq correlation plot for biological replicates of dexamethasone-treated cells (4h, 1μ.M) transfected with the GBS-flank library. (d) The enhancer activity of blunting (n=tral (n=5) and enhancing (n=5) flank variants was assessed for individually transfected STARR-seq constructs by qPCR. Fold change upon dexamethasone treatment normalized to the activity for the scrambled control plasmid is shown as horizontal line for the mean of each activity group and as dot for each individual construct.

Our previous work has shown that the sequence directly flanking GBSs can modulate DNA shape and GR activity [3]. For a parallelized and thorough analysis of sequence variants flanking a GBS, we generated STARR-seq libraries for two GBS variants, we previously named Cgt and Sgk, that showed a strong influence of flanking nucleotides on activity [3]. Specifically, we generated libraries using designed synthetic sequences (synSTARR-seq) containing a GBS with five consecutive randomized nucleotides directly flanking the imperfect half site (Fig. 1A, S2A). Next, we transfected the GBS flank libraries into U2OS-GR cells to determine the activity of each of the 1024 flank variants present in the library. We performed three biological replicates for each condition and found that the results were highly reproducible (r ≥ 0.91 for vehicle treated cells, r ≥ 0.98 for dex treated cells; Fig. 1C, S1B-E). Notably, we retain duplicate reads in our analysis, which is essential to get quantitative information for individual sequence variants of the library. To calculate the activity for each flank variant, we used DESeq2 [28] to compare the RNA-seq read number between dex- and vehicle (ethanol) treated cells (Fig 1A). This resulted in the identification of 189 flank variants with significantly higher activity (enhancing flanks), 125 flank variants with significantly lower activity (blunting flanks) and 710 flank variants that did not induce significant changes in activity neutral flanks). To test the accuracy of the synSTARR-seq data, we cloned 5 flank variants from each activity group (enhancing, blunting and neutral) and assayed the activity of each variant individually by qPCR. Consistent with what we observed for the synSTARR library, the activity of blunting flanks was significantly lower than for the neutral flanks whereas the activity of the enhancing flanks was significantly higher (Fig. 1D). Notably, all flank variants tested were activated upon dex treatment ranging from 2.1 to 15.3 fold (627% higher) depending on the sequence of the flank. Together, our results show that the synSTARR-seq assay produces reproducible and quantitative information and can be used for a high-throughput analysis of the effect of the flanking sequence on GBS activity.

### SynSTARR-seq to assay the effect of flanking nucleotides

To assess how the sequence composition of the flanking region influences GBS activity, we ranked the flank variants by their activity and used a color chart representation to plot the sequence at each position for the Cgt (Fig. 2A) and Sgk GBS (Fig. S2A), respectively. In addition, we generated consensus sequence motifs for the significantly enhancing and blunting variants (Fig. 2B, S2B). Notably, these consensus sequence motifs treat each sequence equally and do not take the quantitative information regarding the activity of each sequence into account. To take advantage of the quantitative information provided by the synSTARR-seq assay, we used *k*pLogo [29], which uses the fold change as weight for each sequence variant, and statistically evaluates the enrichment/depletion of specific nucleotides at each position. The resulting probability logo can be interpreted as an activity logo that visualizes for each position which nucleotides are associated with either higher (letters above the coordinates) or lower (below the coordinates) GBS activity (Fig. 2C, S2C). The activity logo, consensus motifs and color chart highlight several sequence features for enhancing and blunting flank variants. For example, high activity is associated with a T at position 8 for both the Cgt and Sgk GBS, which matches what we found previously when we studied the activity of endogenous GR-bound regions [3]. In addition, the most active flank variants preferentially have an A at position 9 followed by a C at position 10 (Fig. 2A, S2A). To validate that this “TAC” signature results in high activity, we shuffled the sequence to either TCA or CAT and found that this indeed resulted in markedly lower activity (Fig. 2D). For blunting flank variants, we observed a preference for an A at position 8 and a bias against having a C at position 10 (Fig. 2A,C, S2A,C). However, altogether we find that the consensus motifs for enhancing and blunting flanks only have low information content and that a broad spectrum of distinct sequences can enhance or blunt the activity of the adjacent GBS (Fig. 2B, S2B).

**Figure 2.**
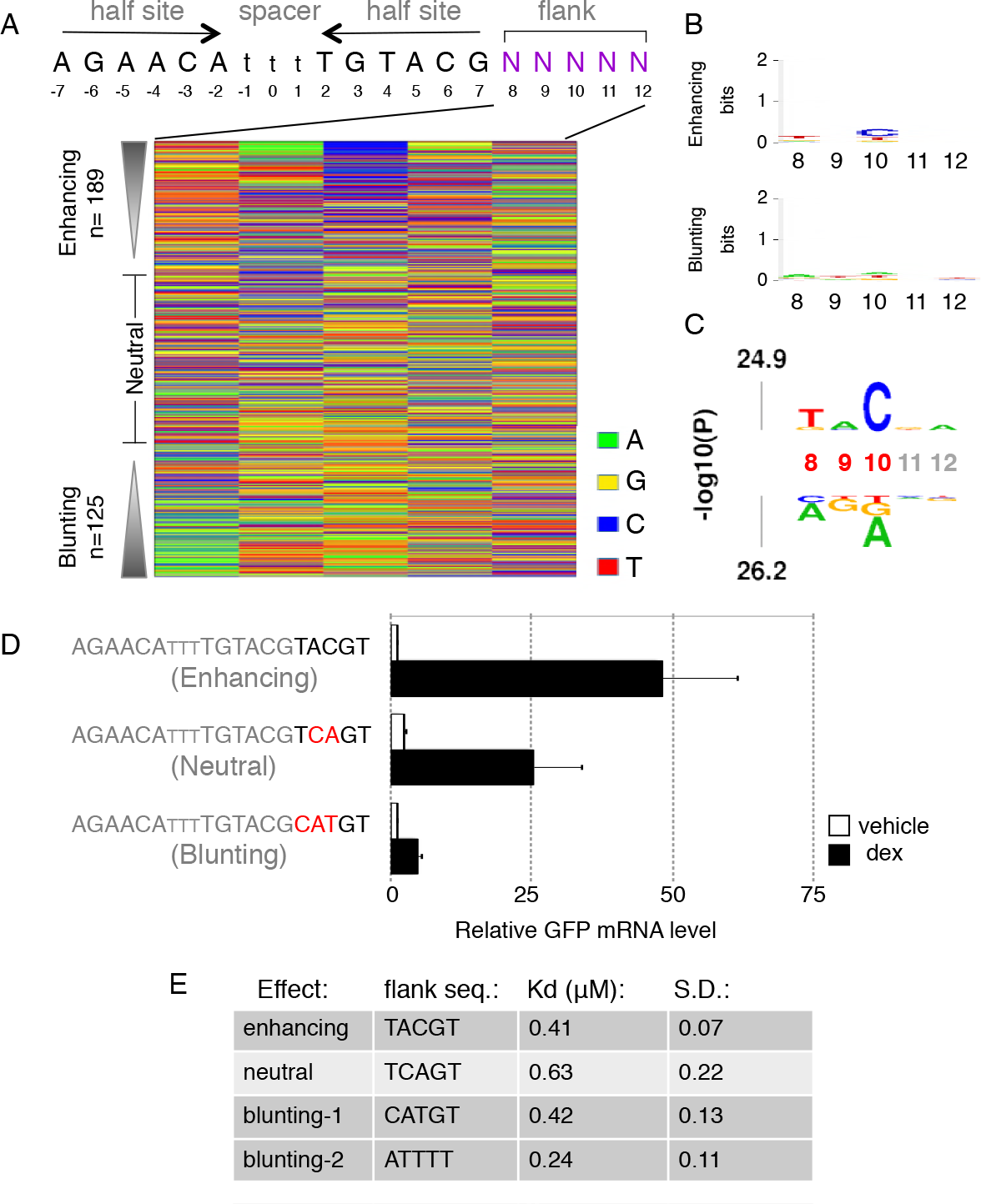
Analysis of the GBS flank library. (a) Color chart summarizing the sequence at each variable position for flank variants ranked by their fold change in response to hormone treatment. (b) Consensus motif for (top) significantly (adjusted p-value < 0.01) enhancing and (bottom) blunting flank variants. (c) *Kp*Logo probability logo (activity logo) for flank variants depicting the p-values from Mann-Whitney U tests of whether GBS variants with a specific nucleotide at a given position are more (displayed above number indicating nucleotide position) or less (displayed below number indicating nucleotide position) active than other GBS variants. Positions with significant nucleotides (p < 0.001) are highlighted (red coordinates). (d) Transcriptional activity of STARR-seq reporters containing candidate flank variants as indicated. Relative RNA levels ± S.E.M. are shown for cells treated with ethanol vehicle and for cells treated overnight with 1 μM dexamethasone (n ≥ 3). (e) Table of EMSA-derived DNA-binding constants (Kd) for flank variants as indicated ± S.D. (n≥3).

Our previous work [3] indicates that DNA shape can influence GR activity downstream of binding. Consistent with this notion, we measured similar Kd values for flanks variants from the different activity classes (Fig. 2E). These findings are also in agreement with published work showing that the nucleotides directly flanking GBSs have little effect on GR affinity [30]. To examine if the flank effects might be explained by differences in DNA shape, we calculated the predicted minor groove width [31] for enhancing and blunting flank variants (Fig. 3A, S2D). Consistent with a role for DNA shape in modulating GR activity, we found shape characteristics that differ between enhancing and blunting flanks. For blunting flanks of the Cgt GBS, we observed a wider minor groove at position 6, and to a lesser degree at position 7 when compared to enhancing flanks Fig. 3A, S3A). In addition, blunting flanks for the Cgt GBS have a narrower minor groove than enhancing flanks for positions 8-12 (Fig. 3A, S3A), a region with several non-specific minor groove contacts with the C-terminal end of the DNA binding domain of GR [5]. For the Sgk GBS library, we find similar shape characteristics associated with blunting flanks with a wider minor groove at position 6 and a more narrow minor groove for positions 8-12 (Fig. S2D, S3B). DNA-shape- based hierarchical clustering recapitulates these characteristics in cluster 4, containing many more blunting flanks than any of the other clusters, for both the Cgt and Sgk GBS flank libraries (Fig. 3B,C, S2E,G). Of note, the consensus motifs for cluster 4 and for the other shape clusters have only low information content (Fig. 3D, S2F) indicating that distinct sequences can give rise to similar shape characteristics with shared effects on the activity of the adjacent GBS.

**Figure 3.**
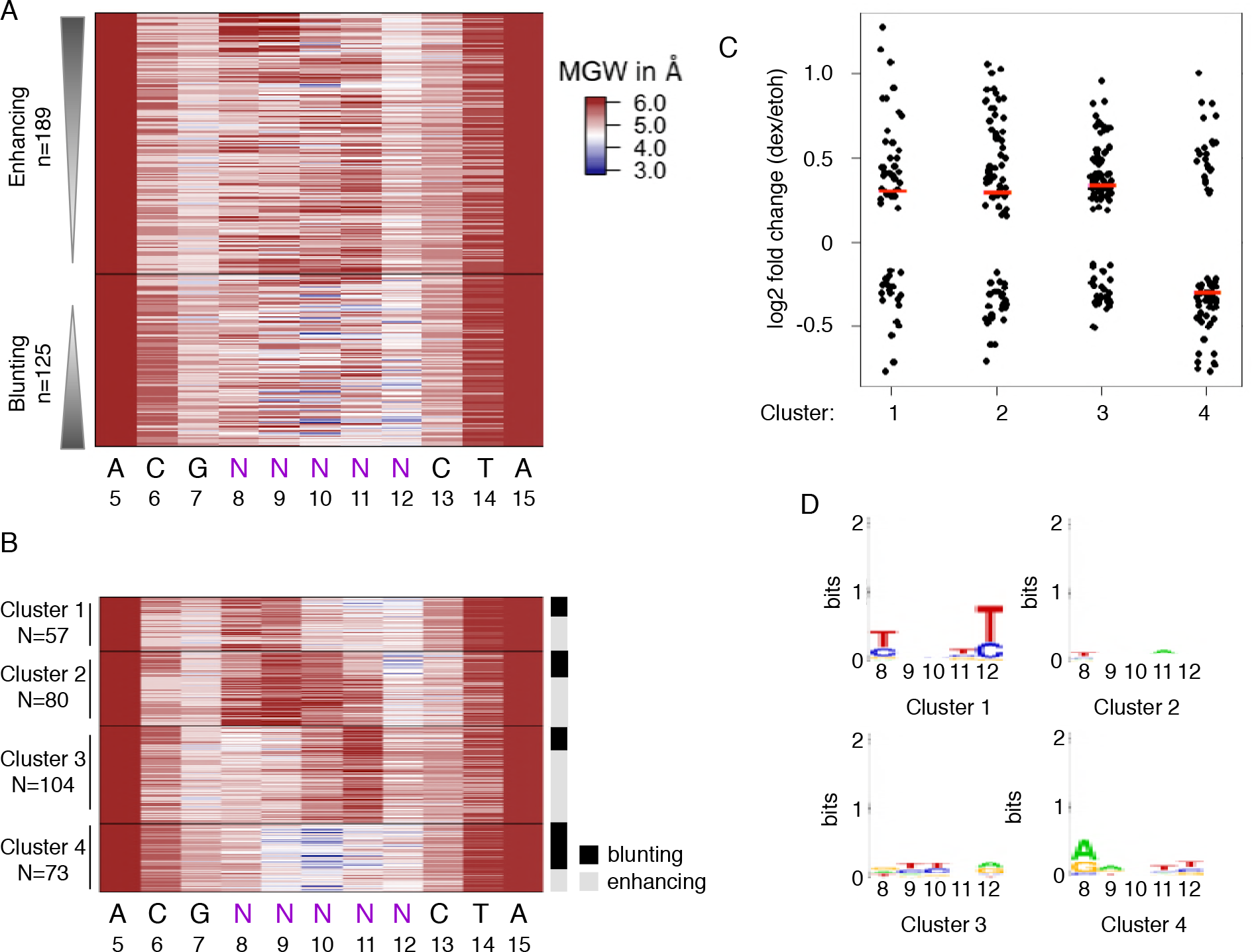
Predicted DNA shape for enhancing and blunting flank variants. (a) Predicted minor groove width (MGW) for significant enhancing and blunting flank variants of the Cgt GBS library ranked by their fold change in response to hormone treatment. (b) K-means clustering based on MGW for significantly enhancing and blunting flank variants. Right side: activating and blunting variants are highlighted in grey and black respectively. (c) Log2 fold change upon dexamethasone treatment for each cluster as indicated. The synSTARR-seq activity for individual sequences is shown as black dots, the median for each cluster as a horizontal red line. (d) Consensus sequence motif for clusters as indicated.

Together, these synSTARR-seq experiments uncover how sequence variation in the flanking region of the GBS influences activity and point at a role for DNA shape in modulating GBS activity.

### SynSTARR-seq to assay the effect of variation within the core GBS

We next generated an additional synSTARR-seq library to study the effect of variation within the 15bp core sequence. This library contains a fixed GBS half site followed by eight consecutive randomized nucleotides (Fig. 4A). The library, containing over 65.000 variants, was transfected into U2OS-GR cells and the read count for each variant was determined both in the presence and absence of hormone treatment. Compared to the flank library, we observed a lower correlation between experiments, especially for variants with a low read count, which were removed before further analysis (Fig. S5). Next, we analyzed data from three biological replicates to determine the activity of variants in the library (Fig. 4B). To validate the measured activities, we cloned 4 sequences that repress, 4 that show a weak activation (log2 fold change <2) and 8 strongly activating GBS variants. Consistent with the results from our screen, the three groups showed distinct levels of activity Fig 4B,C). However, for the group of repressed GBS variants we did not recapitulate the observed repression in our screen Fig 4C), indicating that these variants might behave differently in isolation or alternatively, that the repression might be a consequence of issues with data normalization. Notably, a lack of GR-dependent transcriptional repression was also reported in another study using the STARR-seq approach to study the regulatory activity of GR-bound genomic regions [25] indicating that GR might not be able to repress transcription in the STARR-seq context.

**Figure 4.**
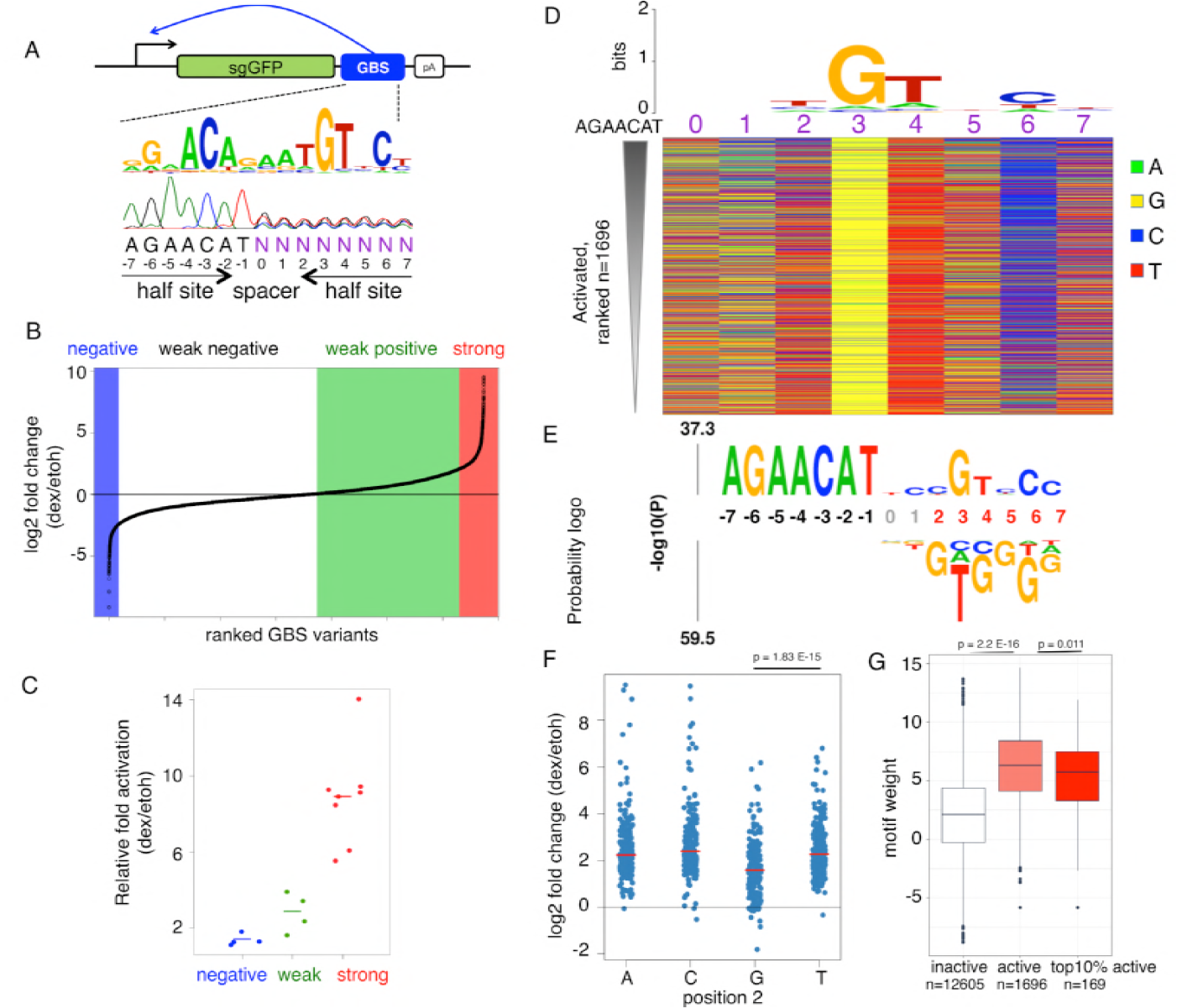
Analysis of the GBS half site library. (a) SynSTARR-seq reporter setup using a synthetic library containing 65536 candidate GR Binding Sequence (GBS) variants (half site library with 8 variable positions N). (b) Candidate GBS variants were ranked by their fold change in expression in response to hormone treatment (4 h, 1 μM dex). Only sequences with a mean read count > 100 across all replicates (n=3) for both dex and ethanol vehicle treated cells are shown. Repressed (log2 FC < -2), weakly active (0 < log2 FC <2) and activated GBS variants (log2 FC ≥ 2) are highlighted by a blue, green and red background respectively. (c) The enhancer activity of negative (n=4), weak (n=4) and strong (n=8) GBS variants was assessed by qPCR for individually transfected STARR-seq constructs. Fold change upon dexamethasone treatment normalized to the activity for the scrambled control plasmid is shown. Horizontal line shows the mean for each activity group; dots the values for individual constructs. (d) Top: Consensus motif and below a color chart summarizing the sequence at each variable position for each significantly activated GBS variant (adjusted p-value < 0.01) ranked by their fold change in response to dex treatment. (e) μpLogo probability logo (activity logo) for half site variants depicting the p-values from Mann-Whitney U tests of whether GBS variants with a specific nucleotide at a given position are more (displayed above number indicating nucleotide position) or less (displayed below number indicating nucleotide position) active than other GBS variants. Positions with significant nucleotides (p < 0.001) are highlighted in red, fixed positions in black. (f) Log2 fold change upon dexamethasone treatment for GBS-like variants with either an A, C, G or T at position 2 (exact match to AGAACATnnXGTnCn, with X either A,C,G or T). Data for individual sequences are shown as blue dots. Horizontal red lines show the median for each group. p-values were calculated using a Student’s t-test. (g) Boxplot of the motif weight (using the truncated 15nt long M00205 motif from Transfac) for inactive (-0.5≤ log2 fold change ≤0.5; white), active (light red) and the top 10% active (dark red) GBS variants. p-values were calculated using a Student’s t-test.

Given that the observed repression was not reproducible, we concentrated our analysis on 1696 sequences that facilitated significant GR-dependent transcriptional activation. Consistent with activation, we found that the consensus motif for activating sequence variants recapitulates the known GR consensus sequence with the second half site 3-bp downstream of the fixed first half site of our library (Fig. 4D). Accordingly, the GBS motif weight, which serves as a proxy for DNA binding affinity, is higher for activating sequences when compared to sequences that did not respond to hormone treatment Fig. 4G). However, the score for the top 10% most active sequences was not higher than for all active variants Fig. 4G), arguing that higher affinity does not drive the high levels of activation. As expected and consistent with the GR consensus motif, the color chart (Fig. 4D) and activity logo (Fig. 4E) highlight a strong preference for a G at position 3 and accordingly GBS activity is significantly lower for variants with a nucleotide other than G at this position (Fig. S7A). The activity logo also highlights that a G at position 2 is associated with lower activity (Fig. 4 E,F).

Previous studies have shown that the sequence of the spacer can modulate GBS activity [4, 5]. Therefore, we compared the activity of all 16 spacer variants in our library that match the GBS consensus for the second half site at the key positions 3, 4 and 6 (Fig. S6A). In line with a role for the spacer in modulating transcriptional output, we find significant differences between the spacer variants (Fig. S6B). For example, the activity for variants with an AC spacer is significantly higher than for most other spacer variants Fig S6B) whereas the activity for GT variants is significantly lower (p.adj < 0.01) than either AA, AC or TC variants (Fig S6B).

Unexpectedly, the activity logo and top of the color chart indicated a high activity for variants with a C at position 2 Fig. 4D,E), instead of a T usually observed in the GR consensus motif and from *in vitro* experiments studying the effect of DNA sequence on GR DNA binding affinity [30]. A careful examination of the sequence composition of the most active variants also revealed a preference for TC at the preceding positions within the spacer (Fig. 4E, 5A). To test if the high activity for sequences with a C at position 2 depends on the nucleotide composition of the preceding nucleotides, we changed them to GG and found that this resulted in a marked reduction in GR-dependent activation (Fig 5B, S8A). In addition, we compared the activity between variants with a T or a C at position 2. The activity was higher for the C variant when preceded by TC. However, when we changed the preceding nucleotides to GG the activation was stronger for the T than the C variant (Fig 5B, S8A). These experiments indicated that the high activity for the C variant depends on the preceding nucleotides.

**Figure 5.**
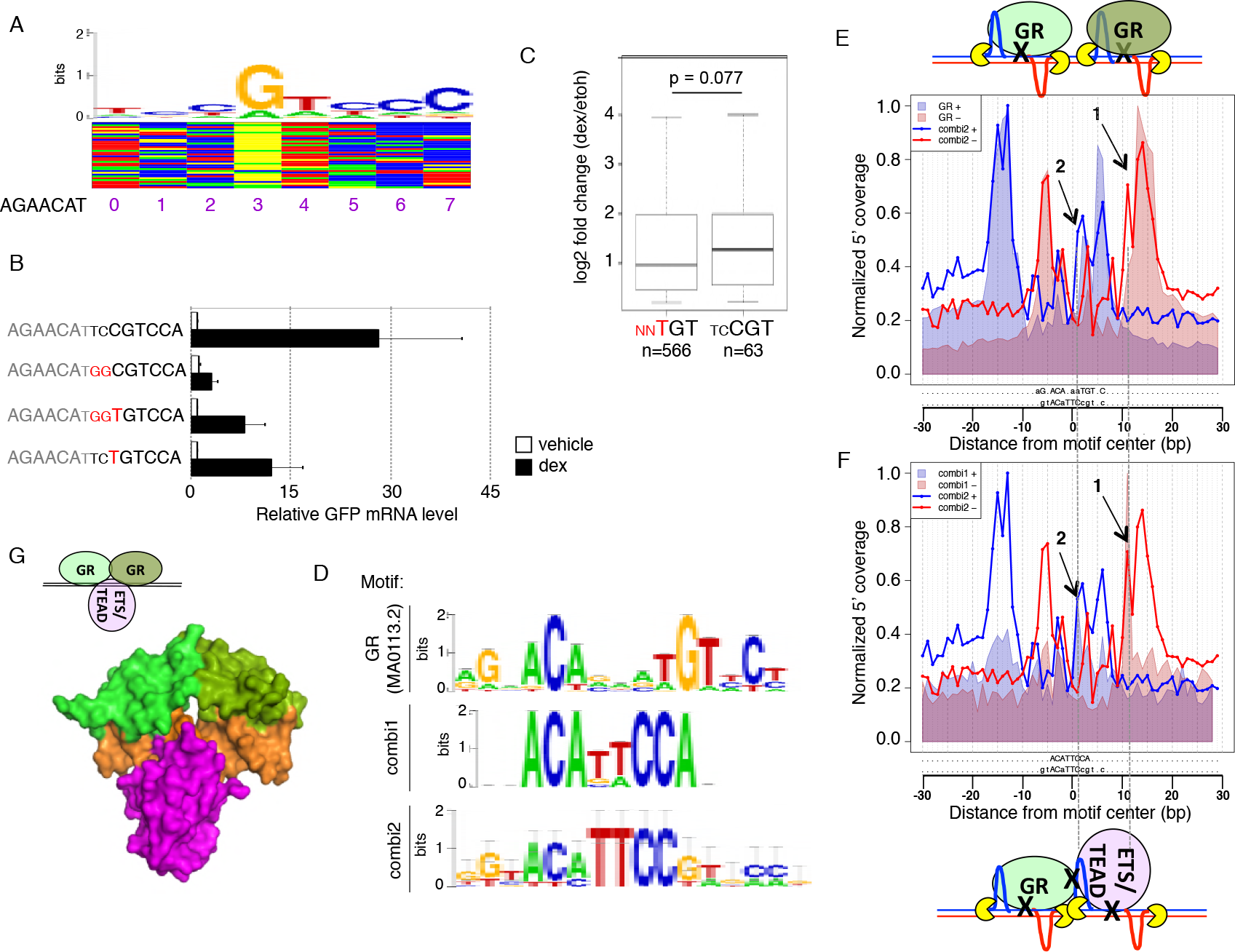
Identification and characterization of the combi2 motif. (a) Color chart for the top activated GBS variants and above the consensus motif for the 25 most active sequences (b) Transcriptional activity of STARR-seq reporters containing candidate GBS variants as indicated. Relative RNA levels ± S.E.M. are shown for cells treated with ethanol vehicle and for cells treated overnight with 1 μM dexamethasone (n = 3). (c) Boxplot of the log2 fold change upon treatment for 4 h with 1 μM dexamethasone for genes with a ChIP-seq peak in the region ±20 kb around the TSS containing either a conventional GBS match (M00205; p value <0.0001) or a combi2-like sequence (combi2 motif; p value < 0.0001). Center lines show the median. p-value was calculated using the Wilcoxon rank sum test. (d) Motif logo representing the positional weight matrices for the canonical GBS (JASPAR MA0113.2), combi1 and combi2 motif. (e) Alignment of the ChIP-exo footprint profiles for the combi2 and the conventional GBS motif. Arrows 1 and 2: Additional 5’ coverage for the combi2 motif that does not match the conventional GBS footprint. (f) Alignment of the ChIP-exo footprint profiles for the combi1 and combi2 motif. Arrows 1 and 2: Additional 5’ coverage for the combi2 motif when compared to the GBS footprint aligns with signal for the combi1 footprint. (g) Structural alignment of combined binding of a GR dimer (green) and ETS1 (purple, middle: PDB 1K79) at the combi2 sequence (orange).

Interestingly, the most active variants resemble the sequence composition of the “combi” motif we identified previously [32]. The combi motif contains only a single GR half site followed by TTCC and we found evidence that GR binds this sequence as a monomer in conjunction with a partnering protein [32]. In contrast to the combi motif, the most active variants from our screen (named “combi2”) also contain a recognizable second half site. To gain insight into the mode of GR binding at the combi2 motif, we examined published ChIP-exo data [32]. ChIP-exo is an assay that combines ChIP with a subsequent exonuclease step [33] which results in a base-pair resolution picture of GR binding. The ChIP-exo signal takes the form of sequence-specific peak patterns (footprint profiles), detectable on both strands with the program ExoProfiler [32]. We applied ExoProfiler to scan GR-bound regions with the combi2 motif (Fig. 5D,E, solid lines). As control, we analyzed the footprint profile for the canonical GR consensus motif (Fig. 5D; JASPAR MA0113.2) and recovered peak pairs on the forward and reverse flanks that demarcate the protection provided by each of the monomers of the GR dimer (Fig. 5E, shaded area). The signal for the first half site is essentially the same and a similar pattern is also observed for the second half site, indicating that GR binds as a dimer on regions bearing the combi2 motif, however with additional signal (highlighted with black arrows in Fig. 5E). In addition, we compared the footprint profile between the original combi (Fig. 5D; [32]) and the combi2 motif (Fig. 5F). Again, the position and shape of the peaks are compatible for the first half site but the ChIPexo signal for the second half site looks markedly different. The aforementioned additional signal for the combi2 motif aligns with the position of the second peak pair of the combi motif (Fig. 5F), indicating that the footprint profile for the combi2 motif appears to be a composite of the signal for homodimeric GR binding at canonical GBSs and the signal for monomeric GR binding together with another protein. Our previous work suggests that this partnering protein on combi motif might be Tead or ETS2. The ChIP-exo profile thus points to three alternative binding configurations on combi2: homodimeric GR, monomeric GR binding with Tead/ETS2 or the simultaneous binding of homodimeric GR complex together with Tead/ETS2. Structural modeling suggests that this third mode is possible given the absence of obvious sterical clashes that would prevent this mode of binding Fig. 5G).

To assess if DNA shape could play a role in modulating GBS activity, we calculated the predicted minor groove width for all 1696 significantly activated sequences ranked by activity (Fig S7B). Comparison of the top 20% most active and bottom 20% least active sequence variants highlighted two regions with significant differences. First, consistent with our findings for the flank library, we find that a wider minor groove at positions 6 and 7 correlates with weaker activity (Fig. S7B,C). Second, we find that a narrower minor groove in the spacer position -1 and 0) correlates with weaker activity Fig. S7B,C). As we observed for the flank variants, the different activity classes do not show a distinct sequence signature (Fig. S7B) again arguing that DNA shape might modulate GBS activity.

Together, the findings for our half site library suggest a role for both DNA shape and sequence in tuning the activity of GBS variants. Moreover, our screen uncovered a novel high-activity functional GR binding sequence variant.

### SynSTARR to assay the effect of enhancer sequence composition on noise

Thus far, we analyzed the effect of sequence composition on transcriptional output by analyzing mean expression levels for populations of cells. To test if sequence variation in the enhancer influences cell-to-cell variability in gene expression (noise), we measured GFP levels for individual STARR constructs in single cells (Fig. 6A,B). Cells were transfected with individual constructs along with an mCherry expression construct to remove extrinsic noise, for example caused by differences in transfection efficiency. We first analyzed sequence variants containing a single GBS single GBS group) including known GBSs, two variants matching the combi2 sequence motif and the Cgt GBS with an enhancing flank variant. Consistent with previous findings [5], we found that GBS variants from the single GBS group induced different mean levels of GFP expression. For example, the mean GFP level upon dex treatment was lower for the pal GBS than for the Cgt variant Fig. 6C, orange and red squares). In line with findings by others [16], we observed that transcriptional noise scales with mean expression with lower noise for variants with higher mean expression (Fig. 6C). Next, we assayed two additional groups of sequences with distinct binding sites architectures that both result in more robust GR-dependent activation when compared to single GBS variants (Fig. 6A). The first group contained three instead of one GBS copy triple GBS group) whereas the second group composite group) contains a GBS flanked by a sequence motif for either AP1, ETS1 or SP1, three sequence motifs that can act synergistically with GR [34, 35]. As expected, the mean GFP expression was higher for each member of both the triple GBS and the composite group when compared to the single GBS group (Fig 6A,C). Interestingly, the increase in mean expression we found for the groups of triple GBS and composite enhancers was not accompanied by a decrease in expression noise (Fig. 6C). The high noise to mean expression ratio was especially striking for several triple GBS variants (3xPal, 3xCgt, 3xSgk and 3x Fkbp5-2) but observed in general for each member of the groups of triple and composite enhancers when compared to the single GBS group. Furthermore, enhancer variants with similar mean expression levels (*e.g.* 3xSgk and Ets1+FKBP5-2) can have vastly different noise levels indicating that binding sites architecture can independently tune both mean expression and cell-to-cell variability in gene expression with noisier expression for enhancers with multiple GBSs.

**Figure 6.**
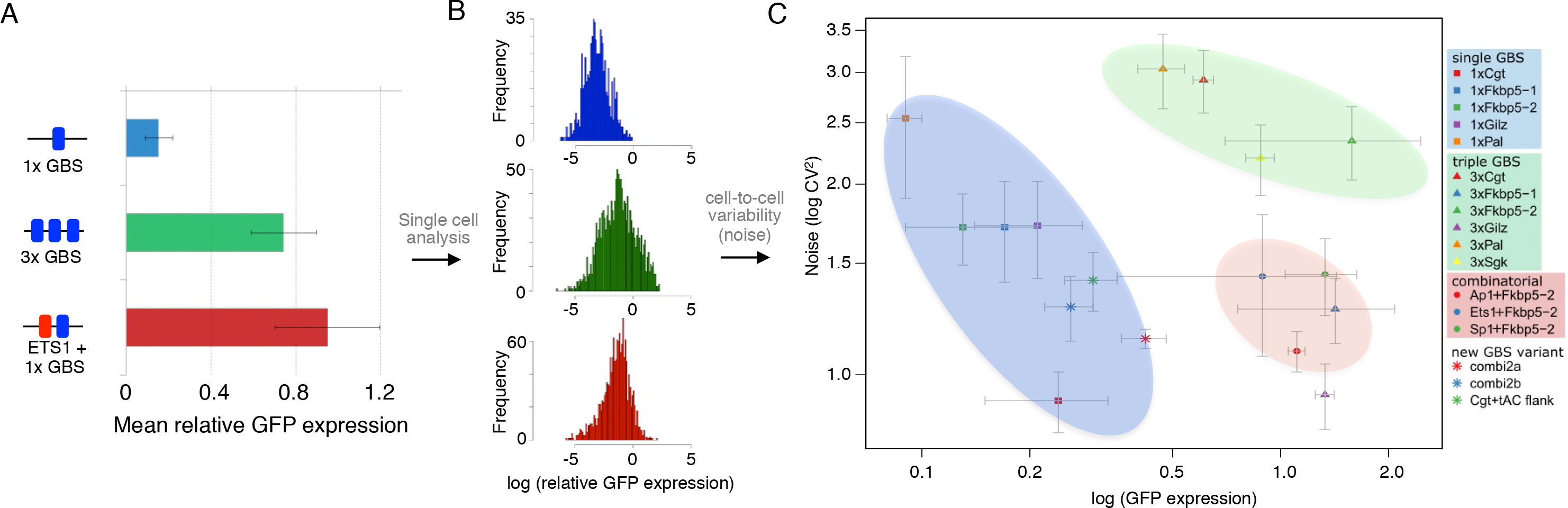
The effect of GBS sequence, number and presence of other TFBS on transcriptional output and noise. (a) Mean GFP expression relative to mCherry of the STARR-seq reporter for cell populations treated overnight with 1 μM dexamethasone with binding site variant as indicated was determined by flow cytometry. (b) The single-cell distribution of GFP expression relative to co-transfected mCherry was determined for each binding site variant as indicated by flow cytometry. The mean and noise for each binding site variant are extracted from these distributions (see Methods). (c) Average and S.D. for mean GFP expression and for noise from three biological replicates. Area with mostly single GBS variants is highlighted with a blue background; Area with three GBSs with a green background and area with background.

## Discussion

In this study, we developed a modified version of the STARR-seq method where we used designed synthetic oligonucleotides to assay how sequence variation within and around the GBS influence GBS activity. This facilitated the thorough and parallelized assessment of 1024 flank variants on GBS activity in a highly reproducible and quantitative fashion (Fig. 1, S1). Similarly, we assessed over 65.000 variants to study how variations in one of the half sites and the spacer influence GBS activity. A key advantage of using designed sequences over the analysis of genomic regions is that variants can be compared in a context where everything is identical except for the sequence of the GR binding site. Notably, the sequence of the binding site is just one of several signals that are integrated at genomic response elements to modulate GR-dependent transcriptional responses. The synSTARR-seq approach can readily be adopted to study how combinations of signals are integrated. For example, principles of combinatorial regulation can be studied using designed sequences for which the GBS is flanked by binding sites for other TFs. Similarly, the assay can be used to investigate the cross-talk between GBS sequence, ligand chemistry, type of core promoter and GR splice isoforms.

Importantly, our findings for the synthetic STARR-seq assay are consistent with GR-dependent regulation of endogenous target genes. Specifically, the nucleotide directly flanking the GBS is preferentially a T for both enhancing flanks in our synSTARR-seq experiments and for the motif we previously found for genomic GR binding sites associated with genes that show the most robust response to GR activation [3]. Moreover, we uncovered a novel functional GR binding sequence variant with high activity, which we called combi2. Consistent with the high activity of the combi2 motif observed in the synSTARR assay, genes with nearby GR-bound peaks matching the combi2 motif were, on average, slightly more activated by GR than genes with peaks matching the consensus motif (Fig. 5C). Other sequence preferences we uncovered for flanks that enhance GBS activity include an A followed by a C at positions 9 and 10 respectively (Fig. 2A,C; S2A,C). One possible explanation for the increased activity is that this sequence generates an additional GR half site or a binding site for another TF. However, the ChIP-exo profile for GBSs flanked by nAC looked essentially the same as the profile for the canonical GBS Fig. S4E), arguing against the binding of an additional factor. Alternatively, the flanking nAC could influence GR’s DNA binding affinity. However, a comprehensive analysis of the effect of sequence variation within and in the regions flanking GR binding sites showed that the flanks essentially do not influence the binding affinity of GR [30]. Accordingly, we found similar Kd values for the AC flank when compared to variants with lower activity Fig. 2E) indicating that the change in activity is not driven by affinity. Together, the synSTARR-seq approach uncovered how sequence variation modulates GR activity, which confirmed previous findings based on a small number of sequences but also provided new insights into mechanisms that modulate GR-dependent regulation of endogenous target genes.

We were surprised to find that the consensus motifs for enhancing and blunting flanks displayed low information content indicating that a broad spectrum of distinct sequences can enhance or blunt the activity of the adjacent GBS (Fig. 2; Fig. S2). However, when looking at DNA shape we found specific shape characteristics for each group (Fig. 3A). This indicates that distinct sequences can induce similar DNA shape characteristics with analogous effects on GBS activity. This finding was corroborated by our analysis of the spacer, which is not directly contacted by GR, yet influences GR activity. Also here we found distinct spacer shape characteristics for the most and least active GBS variants, without a clear sequence signature for each group (Fig. S7B). Furthermore, we trained a model to distinguish between high and low activity GBSs based on either DNA sequence or on predicted minor groove width information. Assessment of the accuracy of the models using ROC curves showed that a single shape parameter, minor groove width, can be used to distinguish quite accurately between blunting and enhancing flanks Fig S9A) and also between the top and bottom 20% active GBS variants (Fig. S9B). Together, our findings which are based on a systematic analysis of many sequence variants are consistent with previous studies based on a small number of binding sites, showing that GR activity can be modulated by DNA shape [3, 4]. Notably, although the role of DNA shape in modulating the affinity of TFs for DNA has been well documented [36–38], we find that DNA shape modulates GR activity without apparent changes in DNA binding affinity (Fig. 2E, [30]). This is consistent with a model where DNA shape acts as an allosteric ligand which induces structural changes in associated TFs which in turn changes the composition and regulatory activity of the complexes formed at the response element [5, 39–41]. Another, not mutually exclusive explanation for flank-dependent modulation of transcriptional output is that flank variants serve as binding sites for other TFs that act additively or synergistically with GR. Further support for the importance of DNA shape comes from the analysis of the conservation of non-coding regions of the genome. This analysis uncovered greater conservation at the level of DNA shape than on the basis of nucleotide sequence indicating that DNA structure may be a better predictor of function than DNA sequence [42]. Accordingly, incorporation of DNA shape characteristics improves *in vivo* prediction of TF binding binding sites [43] and, based on our findings, could also improve the prediction of TF binding site activity.

We also explored if GFP protein expression levels of individual cells can be used to study how enhancer architecture influences cell-to-cell variability in gene expression. A similar approach was used to study how sequence variation of the promoter influences transcriptional noise in yeast [16]. Notably, the only difference between the reporters we assayed is their enhancer sequence, which is downstream of the ORF for the GFP protein. For sequences with related enhancer architectures, we observed transcriptional noise scales with mean expression, such that higher expression levels are associated with lower noise (Fig. 6C). This is consistent with a two-state promoter model where increases in mean expression are driven by an upsurge in transcription burst frequency [44]. Similarly, the estrogen receptor, a hormone receptor closely related to GR, modulates transcription by changing the frequency of transcriptional bursting [12]. When we compare distinct enhancer architectures, we find that expression mean and noise can be uncoupled. Specifically, the noise to mean expression ratio is higher for response elements harboring multiple TF binding sites, indicating that the increase in expression might be accompanied by an increase in the number of transcripts produced during each burst. This finding is consistent with studies in yeast showing that increasing the number of binding sites for GCN4 results in increased expression with relatively high noise levels [16]. Notably, both multiple binding sites for GR and a combination of a GR binding site and a binding site for another TF result in an increased noise to mean expression ratio (Fig. 6). Our results are consistent with a model in which the architecture of the enhancer influences transcriptional burst size and frequency. However, more sophisticated single-cell studies of nascent transcripts are needed for a detailed understanding of the role of enhancer architecture given that our studies are based on the measurement of steady state fluctuations in protein levels. For example, in our experimental approach we cannot rule out that other mechanisms, including differences in RNA stability and translation rates, could contribute to the cell-to-cell variability in expression observed. Nonetheless, our findings argue that differences in enhancer architecture might contribute to gene-specific tuning of expression mean to noise ratios of GR target genes.

## Conclusions

Taken together, we present synSTARR, an approach to measure how designed binding site variants influence transcriptional output and noise. The systematic analysis of sequence variants presented here resulted in the identification of a novel functional GR binding sequence and provides evidence for an important role of DNA shape in tuning GR activity without apparent changes in DNA binding affinity. Our simple approach using designed sequences can be applied to other TFs and can be used to systematically unravel how the interplay between sequence and other signaling inputs at response elements modulate transcriptional output.

## Materials and Methods

### Experimental

#### Plasmids

STARR reporter constructs were generated by digesting the human STARR-seq vector [22] with Sall-HF and AgeI-HF and subsequent insertion of fragments of interest by in-Fusion HD cloning (TaKaRa). All inserts had the following sequence composition: 5’-**TAGAGCATGCACCGG**<u>ACACTCTTTCCCTACACGACGCTCT</u>----*INSERT*----<u>AGATCGGAAGAGCACACGTCTGAACTCCAGTCAC</u>**TCGACGAATTCGGCC**-3’. Sequence homologous to the STARR reporter construct in bold; Sequence for p5 and p7 adaptors underlined. The exact sequence of the insert for each construct used in this study is listed in table S1.

#### Cell lines, transient transfections and luciferase assays

U2OS cells stably transfected with rat GRα (U2OS-GR18) [27] were grown in DMEM supplemented with 5% FBS. Transient transfections were done essentially as described [5] using either lipofectamine and plus reagents (Invitrogen) or using kit V for nucleofections (Lonza).

### Synthetic STARR-seq

<u>Library design and generation:</u> To generate GBS variant libraries, oligos containing degenerate nucleotides N) at defined positions were ordered from IDT as “DNA Ultramer oligonucleotide” sequence listed below). The oligonucleotides were made double stranded using Phusion polymerase (NEB; 98°C for 35 sec, 72°C for 5 min) using the revPrimer (GGCCGAATTCGTCGAGTGAC). The resulting double stranded inserts (25ng) were recombined with 100ng linearized (SalI-HF and AgeI-HF) STARR-seq vector [22] by inFusion cloning in 5 parallel reactions. After pooling the reactions, the DNA was cleaned up using AMPure XP beads (Beckman Coulter), transformed into MegaX DH10B cells (Invitrogen) and plasmid DNA was isolated using a Plasmid Plus Maxi kit (Qiagen). <u>STARR-seq:</u> For STARR-seq experiments, 5 million U2OS-GR18 cells were transfected with 5 μg library-DNA by nucleofection using kit V (Lonza). The next day, cells were treated for 4 h with 1 μM dexamethasone or with 0.1% ethanol as vehicle control. Reverse transcription and amplification of cDNA for subsequence Illumina 50bp paired-end sequencing were done as described [22].

#### Cgt flank library DNA Ultramer oligonucleotide

TAGAGCATGCACCGGACACTCTTTCCCTACACGACGCTCTTCCGATCTCAGCGCAAGAACAtttTGTACGNNNNNCTAGATCGGAAGAGCACACGTCTGAACTCCAGTCACTCGACGAATTCGGCC

#### Sgk flank library DNA Ultramer oligonucleotide

TAGAGCATGCACCGGACACTCTTTCCCTACACGACGCTCTTCCGATCTCAGCGCAAGAACAtttTGTCCGNNNNNCTAGATCGGAAGAGCACACGTCTGAACTCCAGTCACTCGACGAATTCGGCC

#### GBS half site library DNA Ultramer oligonucleotide

TAGAGCATGCACCGGACACTCTTTCCCTACACGACGCTCTTCCGATCTCAGCGAAAGAACAtNNNNNNNNCGTCGCTAGATCGGAAGAGCACACGTCTGAACTCCAGTCACTCGACGAATTCGGCC

### RNA-seq U2OS-GR18 cells (Fig. 5C)

U2OS-GR18 cells were treated for 4h with either 1μM dexamethasone or 0.1% ethanol as vehicle control. RNA was isolated from 1.2 million cells using the RNeasy kit from Qiagen. Sequencing libraries were prepared using the TruSeq RNA library Prep Kit (Illumina). Prior to reverse transcription, poly adenylated RNA was isolated using oligo d T) beads. Paired end 50bp reads from Illumina sequencing were mapped against the human hg19 reference genome using STAR [45] (options: --alignIntronMin 20 --alignIntronMax 500000 -- chimSegmentMin 10 --outFilterMismatchNoverLmax 0.05 --outFilterMatchNmin 10 -- outFilterScoreMinOverLread 0 --outFilterMatchNminOverLread 0 -- outFilterMismatchNmax 10 --outFilterMultimapNmax 5). Differential gene expression between dex and etoh conditions from three biological replicates was calculated with DESeq2 [28], default parameters except betaPrior=FALSE.

### Electrophoretic mobility shift assays

EMSAs were performed as described previously [3] using Cy-5 labeled oligos as listed in Table S2.

### RNA isolation, reverse transcription and qPCR analysis

RNA was isolated from cells treated for either 4 h or overnight with 1 μM dexamethasone or with 0.1% ethanol vehicle. Total RNA was reverse transcribed using gene-specific primers for *GFP* (CAAACTCATCAATGTATCTTATCATG) and *RPL19* (GAGGCCAGTATGTACAGACAAAGTGG) which was used for data normalization. qPCR and data analysis were done as described [5]. Primer pairs for qPCR: hRPL19-fw: ATGTATCACAGCCTGTACCTG, hRPL19rev: TTCTTGGTCTCTTCCTCCTTG, GFP-fw: GGCCAGCTGTTGGGGTGTC, GFP-rev: TTGGGACAACTCCAGTGAAGA.

### Noise-Measurements

For noise measurements, U2OS-GR18 cells were transfected using lipofectamine and plus (Invitrogen) essentially as described [5]. In short: The day before transfection, 40.000 U2OS-GR cells were seeded per well of a 24 well plate. The following day, cells were transfected with individual STARR reporter constructs (20ng/well) along with a SV-40 mCherry expression construct (20ng/well) and empty p6R plasmid (100 ng/ well). Transfected cells were treated overnight with either 1 μM dexamethasone or with 0.1% ethanol vehicle control. Fluorescence intensity was measured using an Accuri C6 flow cytometer (BD Biosciences) and the yellow laser (552nM) and filter 610/20 for mCherry and the deepblue laser (473nM) and filter 510/20 to measure GFP. Gates were set for mCherry and GFP and only cells showing both mCherry and GFP fluorescence were included in the analysis. Relative expression of GFP (GFP/Cherry), from 800-1600 individual dexamethasone-treated cells, was used to calculate mean expression and the standard deviation of cell populations. Mean and standard deviation for noise (CV^2^) and for relative GFP expression were derived from three biological replicates.

### Computational analyses

#### Analysis of synSTARR-seq data

RNA-seq reads were filtered and only sequences exactly matching the insert sequence in length and nucleotide composition were included in the analysis. The number of occurrences for each sequence variants was counted for each experimental condition and differentially expressed sequences were identified using DESeq2 [28] using a p adjusted value <0.01 as cut-off. To fit the dispersion curve to the mean distribution, we used the local smoothed dispersion (DESeqwithfitType=“local”). Notably, each of the constructs of the flank libraries contains a functional GBS. Therefore, flanks that blunt activity will appear repressed after hormone treated because their fraction in the total pool of sequences decreases relative to flank variants with higher activities. For the flank libraries, we obtained information for each sequence variant (1024) in the library. For the half site library, we identified 61.582 out of the 65.536 possible variants present in this library. We found that including sequences with low read coverage resulted in many false positive differentially expressed GBS variants. To avoid this, we only included sequences with a mean read count above 100 across all experiments, leaving us with information for 33.689 sequence variants. The pearson correlation coefficient for replicates was calculated using the ggscatter function of the ggpubr library in R.

Boxplots comparing groups of sequence variants as specified in the figure legends show center lines for the median; box limits indicate the 25th and 75th percentiles; whiskers extend 1.5 times the interquartile range from the 25th and 75th percentiles.

Sequence logos to depict the consensus motif for groups of sequences were generated using WebLogo [46]. The probability logo (activity motif) was generated with *k*pLogo [29] using as input the sequence and fold change dex/etoh) for each variant and the default settings for weighted sequences.

#### Motif weight

The motif weight for each variant was calculated using the RSAT *matrix-scan* program [47, 48]. Specifically, the motif weight was calculated using Transfac motif M00205 truncated to the core 15bp, and a custom background model created with RSAT *create background* program, trained on human open chromatin available at UCSC genome browser (http://genome.ucsc.edu/cgi-bin/hgTrackUi?db=hg19&g=wgEncodeRegDnaseClustered). Boxplots comparing groups of sequence variants show center lines for the median; box limits indicate the 25th and 75th percentiles; whiskers extend 1.5 times the interquartile range from the 25th and 75th percentiles.

#### Comparison of ChIP-seq peak height between combi2 and canonical GBS motif

GR ChIP-seq data sets for U2OS-GR18 cells were downloaded as processed peaks from EBI ArrayExpress (E-MTAB-2731). ChIP-seq peaks in a 40 kb window centered on the transcription start site of differentially expressed genes (RNA-seq data: E-MTAB-6738) were scanned using RSAT *matrix-scan* [47, 48] for the occurrence of either a GBS-match (Transfac matrix M00205, p value cut-off: 10^-4^) or the combi2 matrix we generated (Fig. 5D, p-value cut-off 10^−4^). Next, peaks were grouped by motif match and median peak height was calculated for each group and the p-value comparing both groups was calculated using a Wilcoxon rank-sum test to produce Supplementary Fig. S8B.

#### Comparison of gene regulation

To compare the level of activation between genes with nearby peaks with either a GBS match (Transfac matrix M00205, p value cut-off: 10^−4^) or a combi2 match (motif Fig. 5D, p-value cut-off 10^−4^), we first scanned ChIP-seq peaks (U2OS-GR cells: E-MTAB-2731) in a 40 kb window centered on the transcription start site using all annotated TSSs from Ensembl GRCH37) for motif matches using RSAT *matrix-scan* [47, 48]. Only peaks with an exclusive motif match were retained to generate a boxplot comparing the log2 fold change for genes of each group (RNA-seq data: E-MTAB-2731). Center lines show the median, box limits indicating the 25th and 75th percentiles and whiskers extending 1.5 times the interquartile range from the 25th and 75th percentiles. p-value comparing the log2 fold change for both groups was calculated using a Wilcoxon rank-sum test to produce figure 5C.

#### DNA shape prediction

We used DNAshapeR [31] to predict the minor groove width for sequence variants of interest. Boxplots for individual nucleotide position show center lines for the median; box limits indicate the 25th and 75th percentiles; whiskers extend 1.5 times the interquartile range from the 25th and 75th percentiles. The Wilcoxon rank-sum test was used to calculate the p-values comparing nucleotide position variants between groups. Individual sites were clustered using K-means clustering with k=4 clusters nstart=20 and 100 restarts with the function ‘kmeans’ from the R ‘stats’ package.

#### Classification of GBS activity

To assess classifier performance we generate ROC curves using 10-fold cross-validation. Four different models were tested to classify GBS activity into blunting or enhancing. A mononucleotide model consisting of sequence motifs estimated from relative nucleotide frequencies within the two classes. Class affiliation is predicted with a likelihood ratio test. We also tested a similar model based on dinucleotides. In addition, we tested two random forest (RF) classifiers with 100 trees, based on sequence and shape information. We used the R package “randomForest” for constructing the classifiers [49]. Since RF classifiers are not designed for categorical data, we coded nucleotide sequences using 00 for ‘A’, 01 for ‘C’, 10 for ‘G’, and 11 for ‘T’.

#### ChIP-exo foo tprin t profiles

ChIP-exo footprint profiles were generated using the ExoProfiler package [32] and published ChIP-exo (EBI ArrayExpress E-MTAB-2955) and ChIP-seq (E-MTAB-2956) data for IMR90 cells as input. Peaks were scanned using either the JASPAR MA0113.2 motif [50], the PWM for the combi1 motif [32], the combi2 motif (Fig. 5D) or for the AC flank variant, the motif depicted in figure S4A. Hits were included if the p-value was <10^-4^. Overlay plots for distinct motifs were generated by aligning the profiles on the GBS and normalizing the signal for each motif variant to 1.

#### Structural al/gnment of GR:ETS1 complex

Structural alignment of the GR:ETS1 complex on a combi2 sequence was done as described previously [32] except that both GR dimer halves are retained in the resulting model. In short: A structural model of the DNA hybrid sequence AGAACATTCCGGCACT) was generated using 3D-Dart [51] using the ETS1 structure (PDB entry 1K79) and the GR structure (PDB entry 3G6U). GR and the ETS2 binding motifs were aligned using the CE-align algorithm [52] to the 3D-DART DNA model of the hybrid sequence.

## Data access

Data were deposited in ArrayExpress under the accession numbers: E-MTAB-6738 (RNA-seq U2OS-GR18) and E-MTAB-6737 (synSTARR-seq U2OS-GR18). In addition, we used the previously deposited datasets: E-MTAB-2731 (ChIP-seq U2OS cells), E-MTAB-2955 and E-MTAB-2956 (ChIP-seq and ChIP-exo data IMR90).

## Reviewers access to datasets

STARR-seq data: E-MTAB-6737
Username: Reviewer_E-MTAB-6737
Password: hgeofcho
RNA-seq data: E-MTAB-6738
Username: Reviewer_E-MTAB-6738
Password: cieef7tt

## Funding

This work was supported by the Deutsche Forschungsgemeinschaft [ME4154/1-1 to SS and MJ].

## Acknowledgements

We would like to thank Marcel Jurk for performing the structural alignment presented.

## Author’s contributions

S.S., M.B., M.B., M.T.C., E.E., P.B. and S.H.M. performed and conceived experiments and analyzed the data. S.S., M.T.C., M.V. and S.H.M. designed and supervised the study and wrote the manuscript with input from all authors.

## Supplementary figure legends

**Figure S1.**
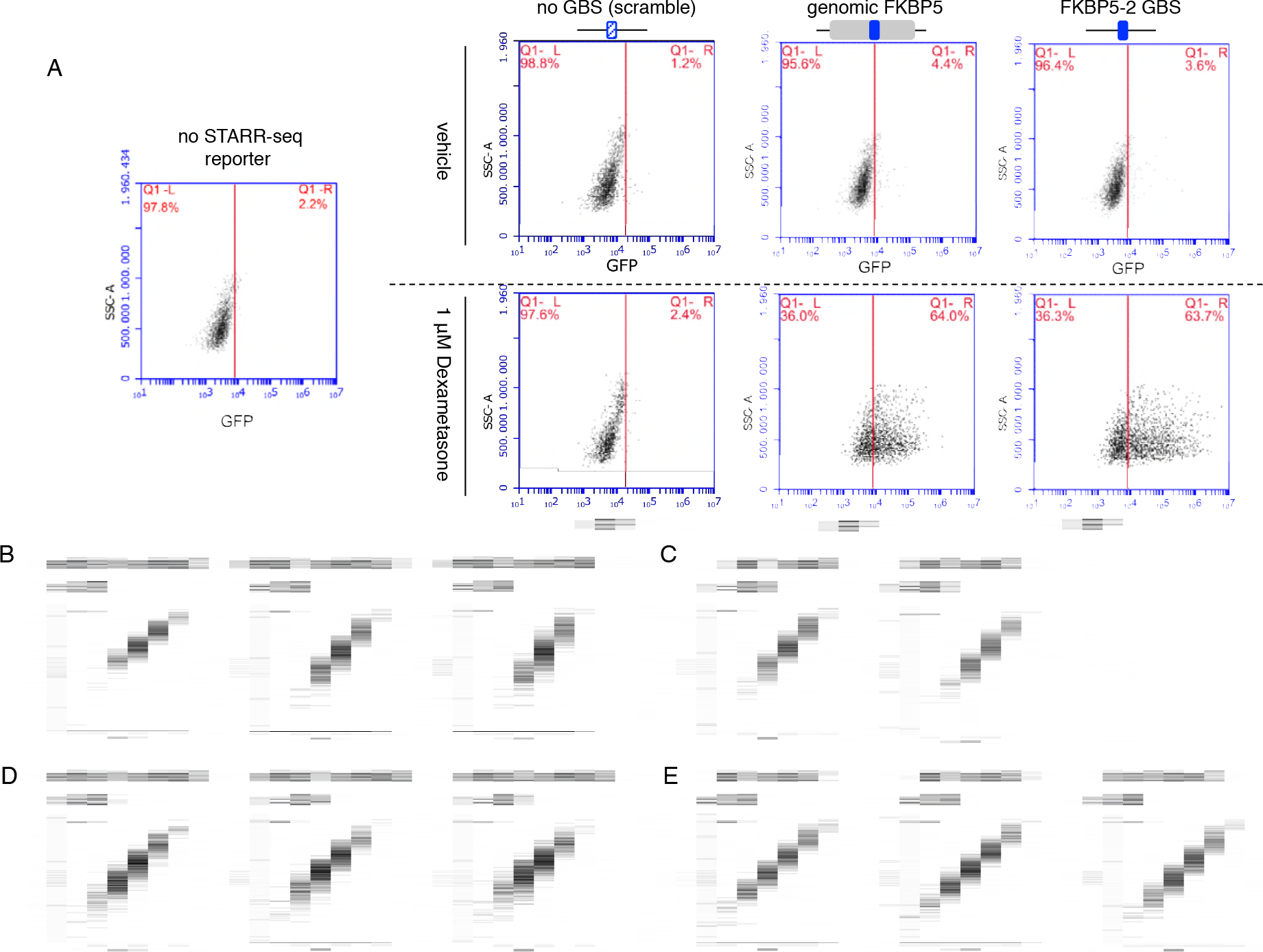
Analysis of individual enhancer variants by flow cytometry and synSTARR-seq reproducibility. (a) Analysis of individual enhancer variants as indicated by flow cytometry showing the side scatter (SSC-A) versus GFP signal for individual mCherry-positive cells. Left: no STARR-seq construct. Right-Top: ethanol, vehicle, treated cells; Right-Bottom: Cells treated overnight with 1μM dexamethasone. Numbers in red indicate the percentage of GFP+ (top right side) and GFP-(top left side) cells respectively. Red vertical line demarcates the threshold for being called GFP+. (b) RNA-seq correlation plots for biological replicates of vehicle-treated cells transfected with the GBS-flank library (Cgt flank library). (c) Same as (b) except for biological replicates of dexamethasone-treated cells (4h 1μM). (d) RNA-seq correlation plots for biological replicates of vehicle-treated cells transfected with the GBS-flank library (Sgk flank library). (e) Same as (d) except for biological replicates of dexamethasone-treated cells (4h 1μM).

**Figure S2.**
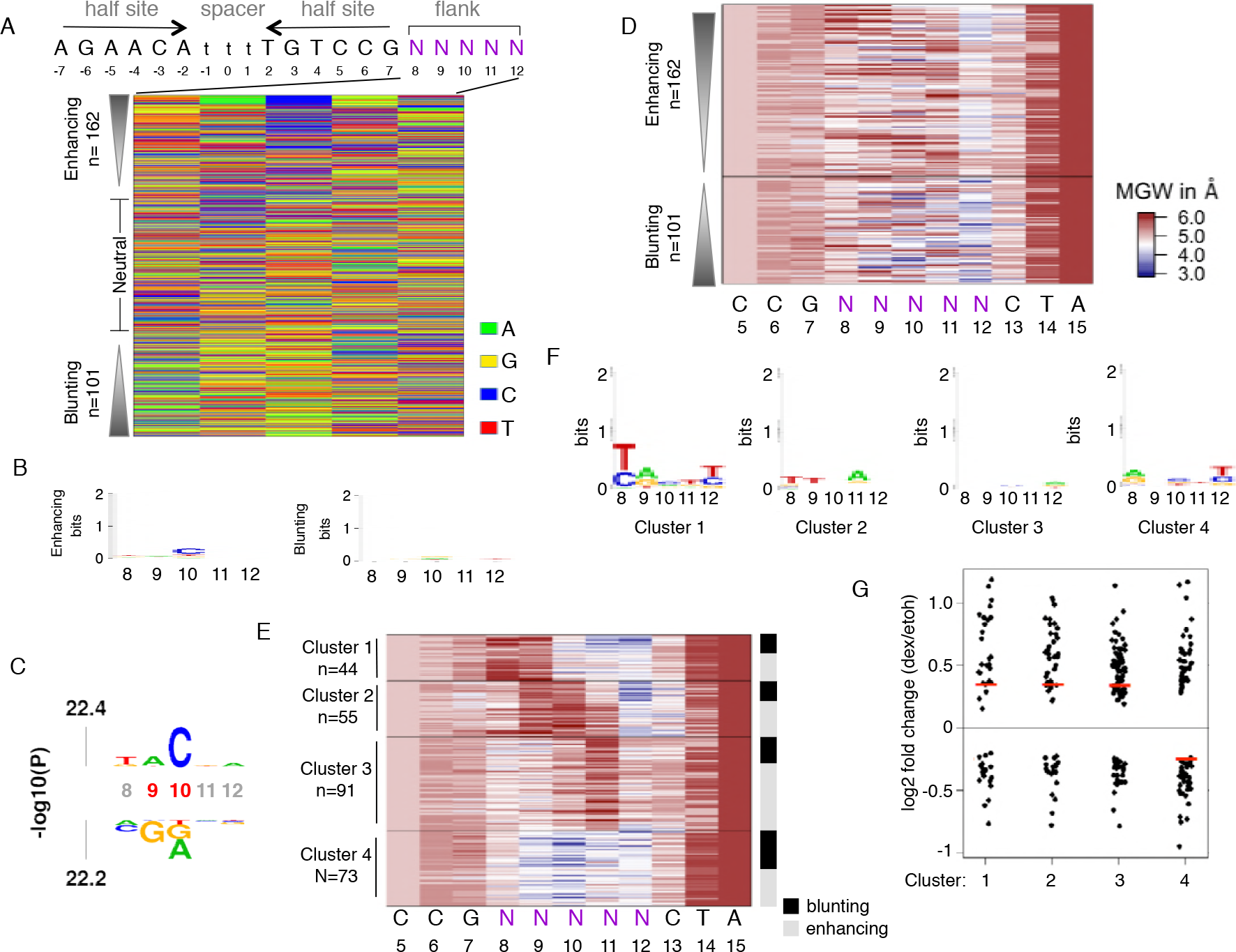
Analysis of the Sgk flank library. (a) Color chart summarizing the sequence at each variable position for flank variants ranked by their fold change in response to hormone treatment. (b) Consensus motif for (left) significantly enhancing and (right) blunting flank variants (c) *k*pLogo probability logo (activity logo) for flank variants depicting the p-values from Mann-Whitney U tests of whether GBS variants with a specific nucleotide at a given position are more (displayed above number indicating nucleotide position) or less (displayed below number indicating nucleotide position) active than other GBS variants. Positions with significant nucleotides (p < 0.001) are highlighted (red coordinates). (d) Predicted minor groove width (MGW) for significant enhancing and blunting flank variants of the Sgk GBS library ranked by their fold change in response to hormone treatment. (e) K-means clustering based on MGW for significantly enhancing and blunting flank variants. Right side: activating and blunting variants are highlighted in grey and black respectively. (e) Consensus sequence motif for clusters as indicated. (g) Log2 fold change upon dexamethasone treatment for each cluster as indicated. The synSTARR-seq activity for individual sequences is shown as black dots, the median for each group as a horizontal red line.

**Figure S3.**
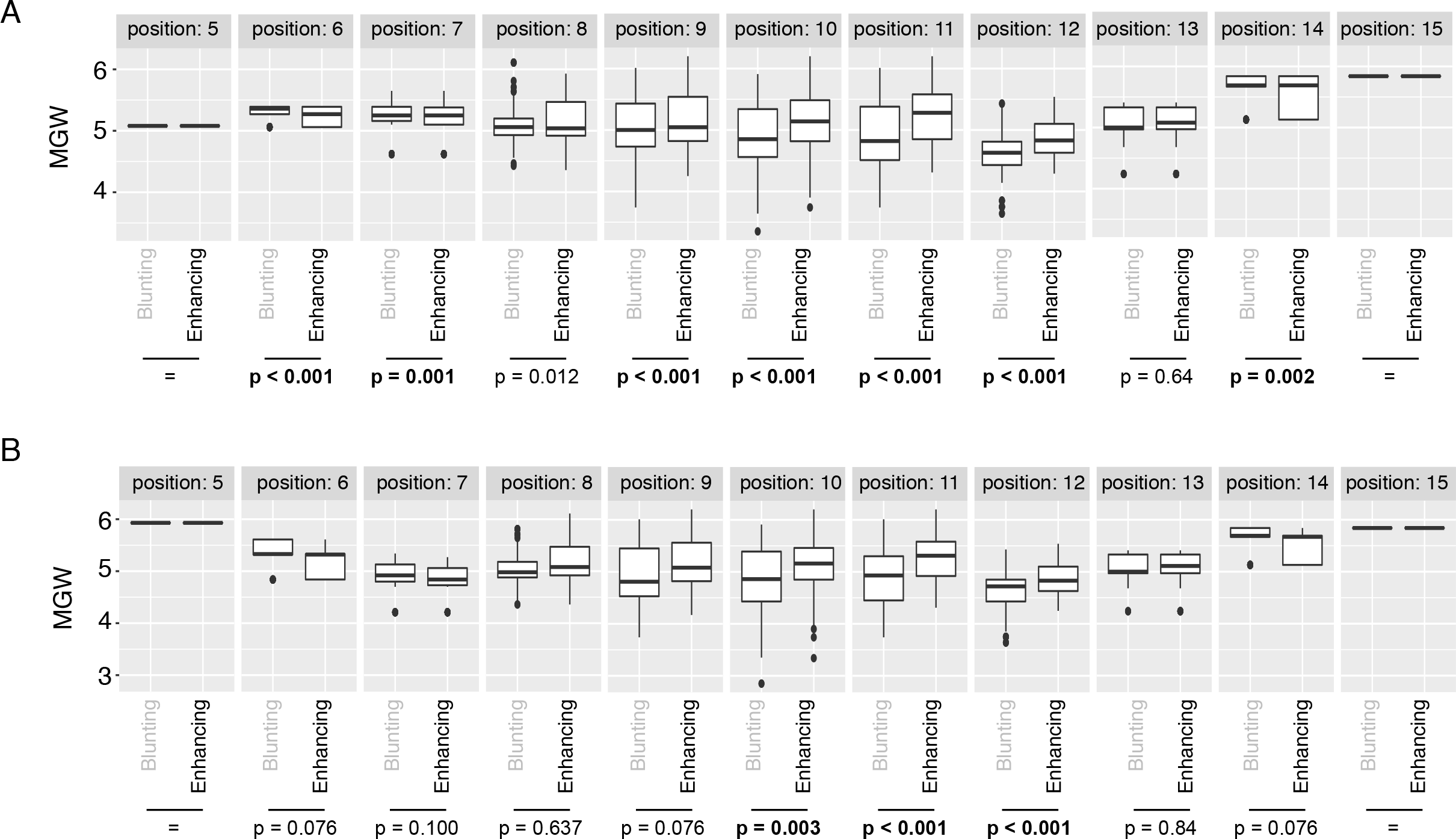
MGW comparison between blunting and enhancing flanks. (a) Minor groove width (MGW) for selected individual bases for significantly blunting (n=189) and significantly enhancing (n=125) flanks for the Cgt library. p-values were calculated using the Wilcoxon rank-sum test. (b) Same as for (a) except for significantly blunting (n=162) and significantly enhancing (n=101) flanks of the Sgk flank library.

**Figure S4.**
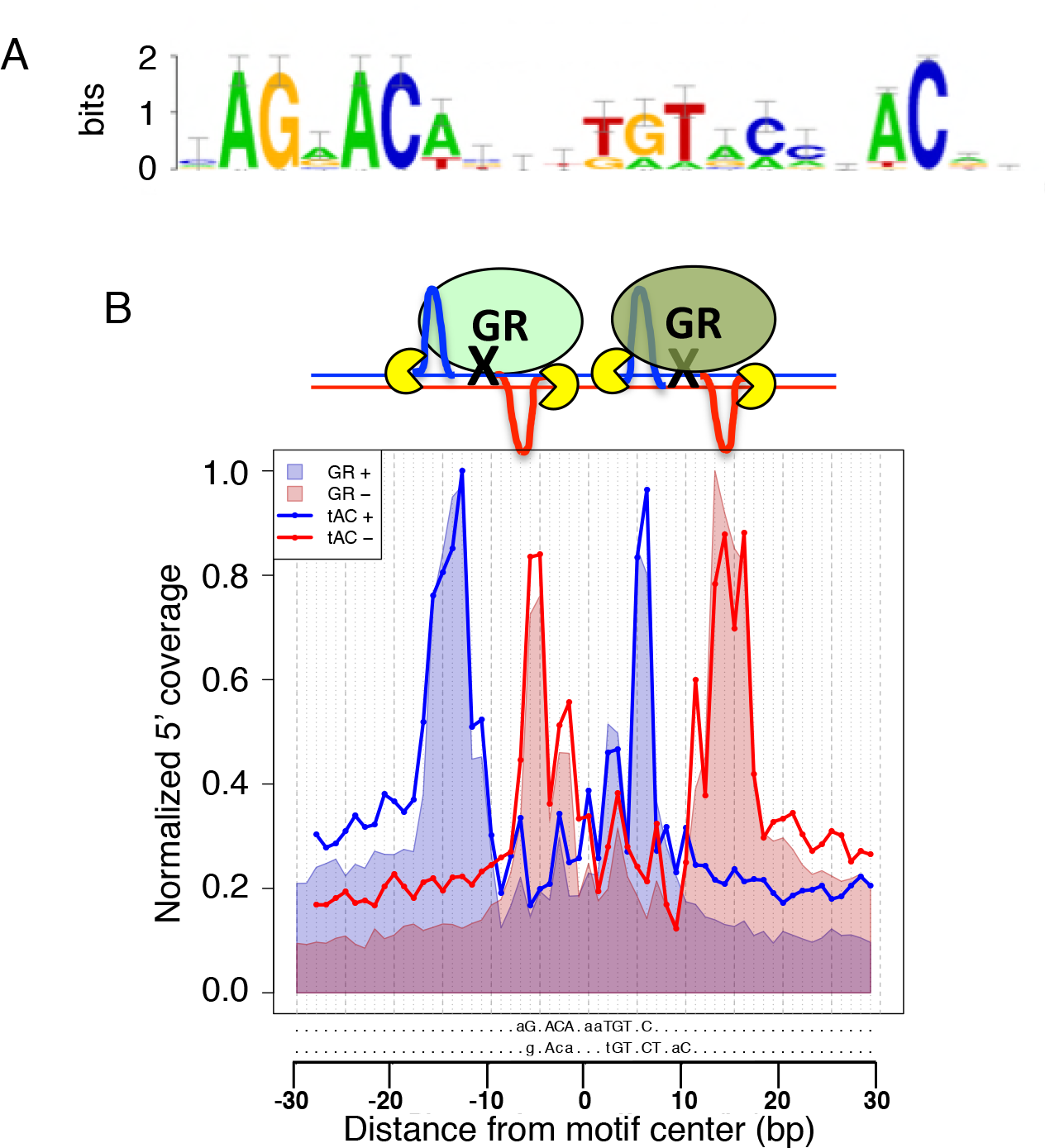
Analysis of the nACnn flank. (a) Motif logo representing the positional weight matrix of highly active flank variants that was used to scan for motif-matches to generate the ChIP-exo footprint profile. (b) Alignment of the ChIP-exo footprint profiles for highly active flank variant matches (p value <0.0001; solid lines: blue: positive strand, red: negative strand) and for the conventional GBS motif (M00205; p value <0.0001; shaded areas; blue: positive strand, red: negative strand).

**Figure S5.**
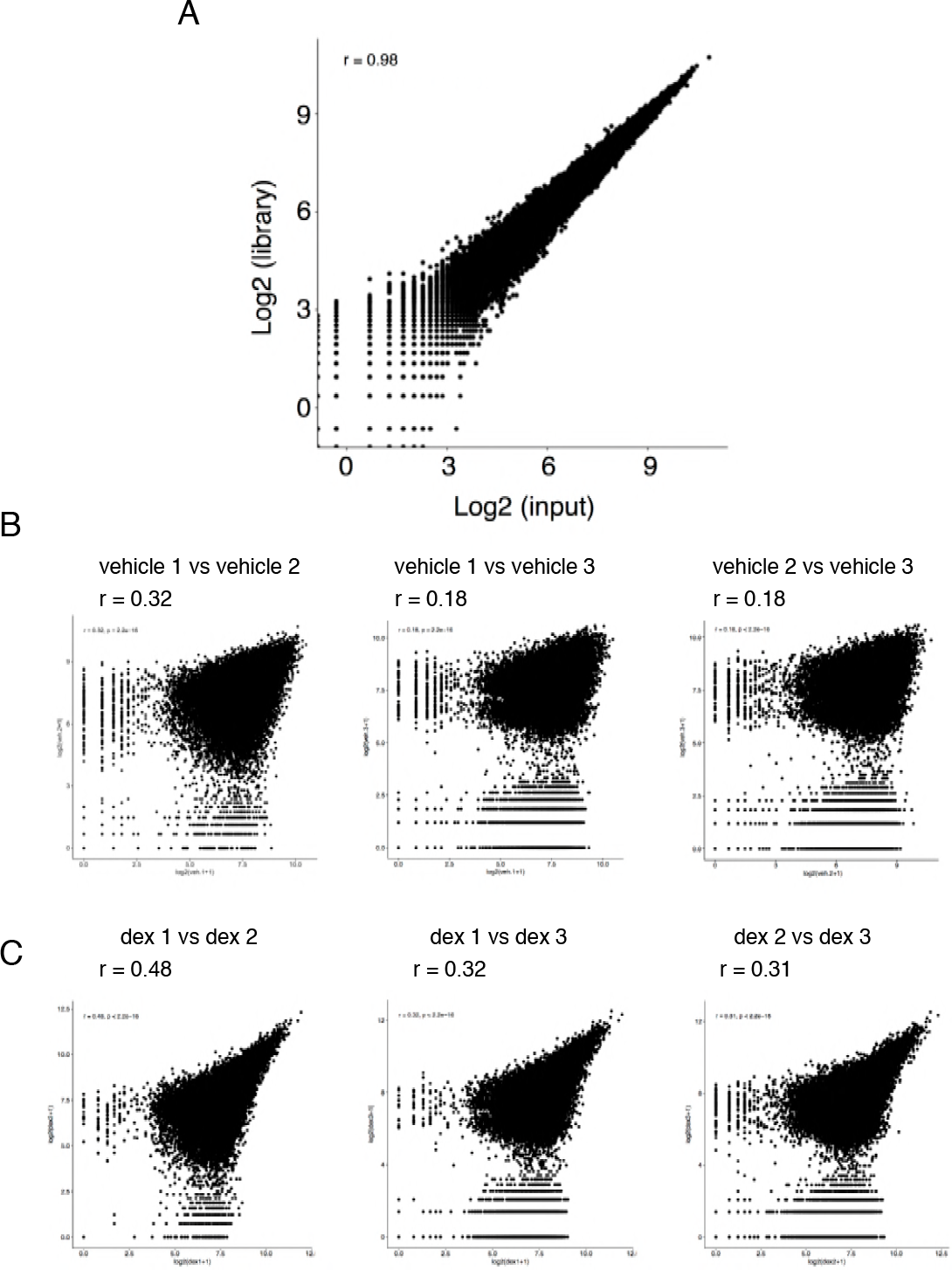
synSTARR-seq reproducibility for the half site library. (a) Correlation plot between input library (library) and the plasmid library isolated from transfected U2OS-GR18 cells (input). (b) RNA-seq correlation plots for biological replicates of vehicle-treated cells. (c) Same as for (b) except for biological replicates of dexamethasone-treated cells (4h 1μM).

**Figure S6.**
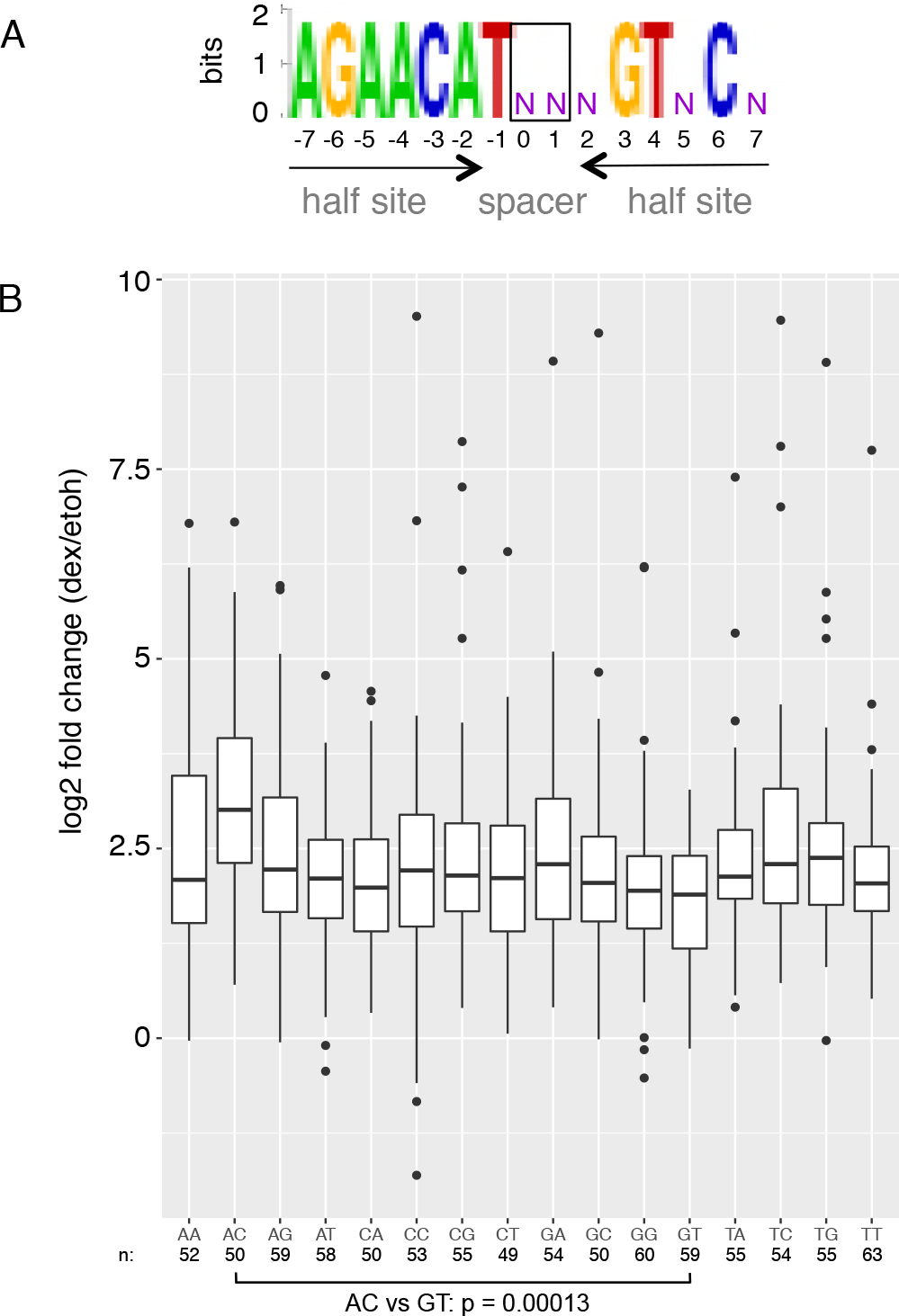
Effect of spacer sequence on GBS activity. (a) Motif logo representing the sequence that was used to scan for GBS-matches in the half site library. Black box highlights the two positions in the spacer whose effect on GBS activity was assayed. (b) Boxplot of the log2 fold change upon treatment for 4 h with 1 μM dexamethasone for GBS matches with spacer variant as indicated. Center lines show the median. The Benjamini-Hochberg corrected p-value for the spacer variants with the most significance difference was calculated using a Student’s t-test.

**Figure S7.**
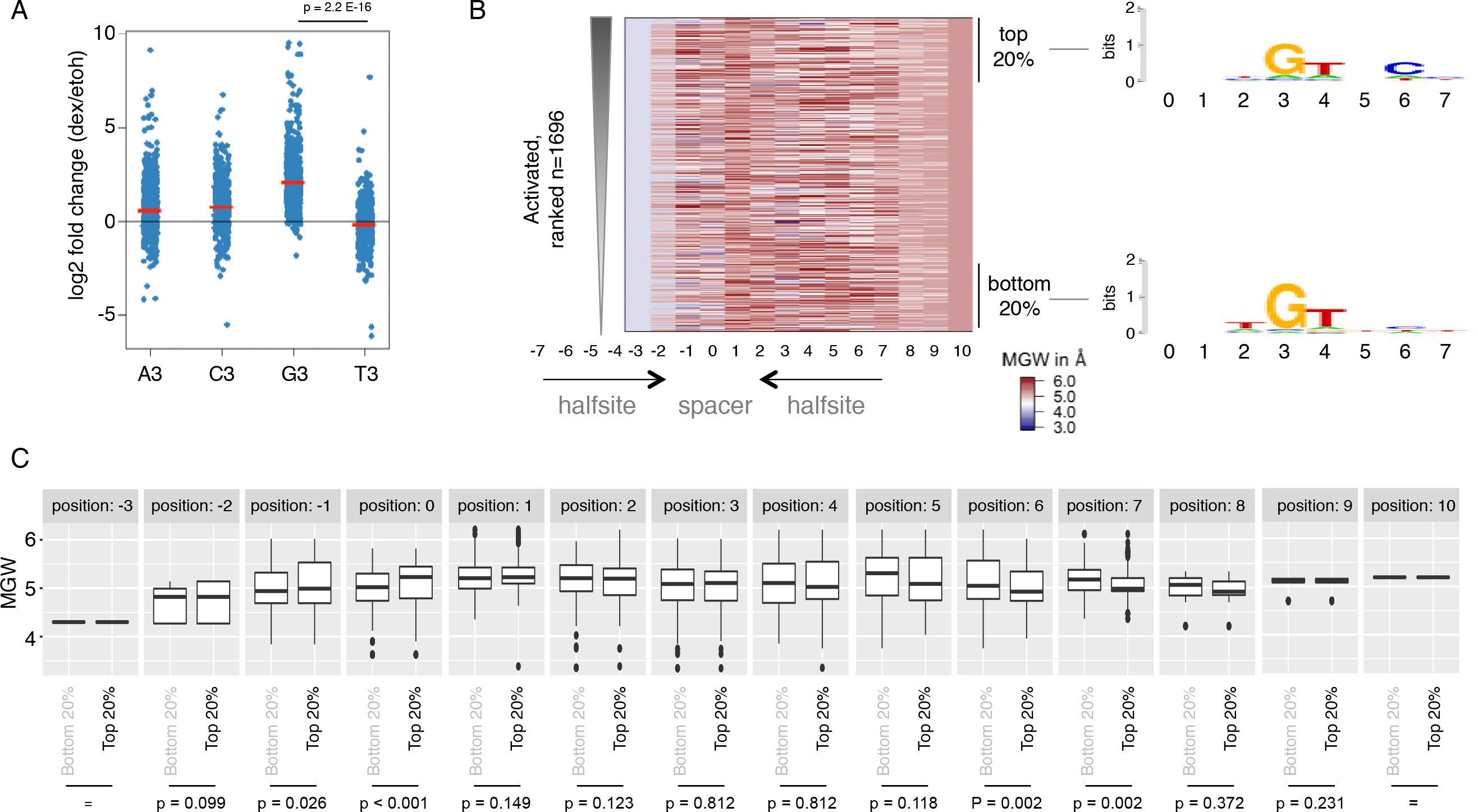
Analysis of the GBS half site library. (a) Log2 fold change upon dexamethasone treatment for active GBS variants with either an A, C, G or T at position 3. Data for individual sequences that match consensus second half site at key positions 4 and 6 (exact match to AGAACATnnnXTnCn, with X either A,C,G or T) are shown as blue dots. Horizontal red lines show the average for each group. p-value was calculated using a Student’s t-test. (b) Left: Minor groove width (MGW) prediction for GBS variants ranked by activity. Right: Consensus motif for top 20% most active and bottom 20% least active GBS variants. (c) MGW for select individual bases comparing the top 20% most active and bottom 20% least active activated GBS variants. p-values were calculated using the Wilcoxon rank-sum test.

**Figure S8.**
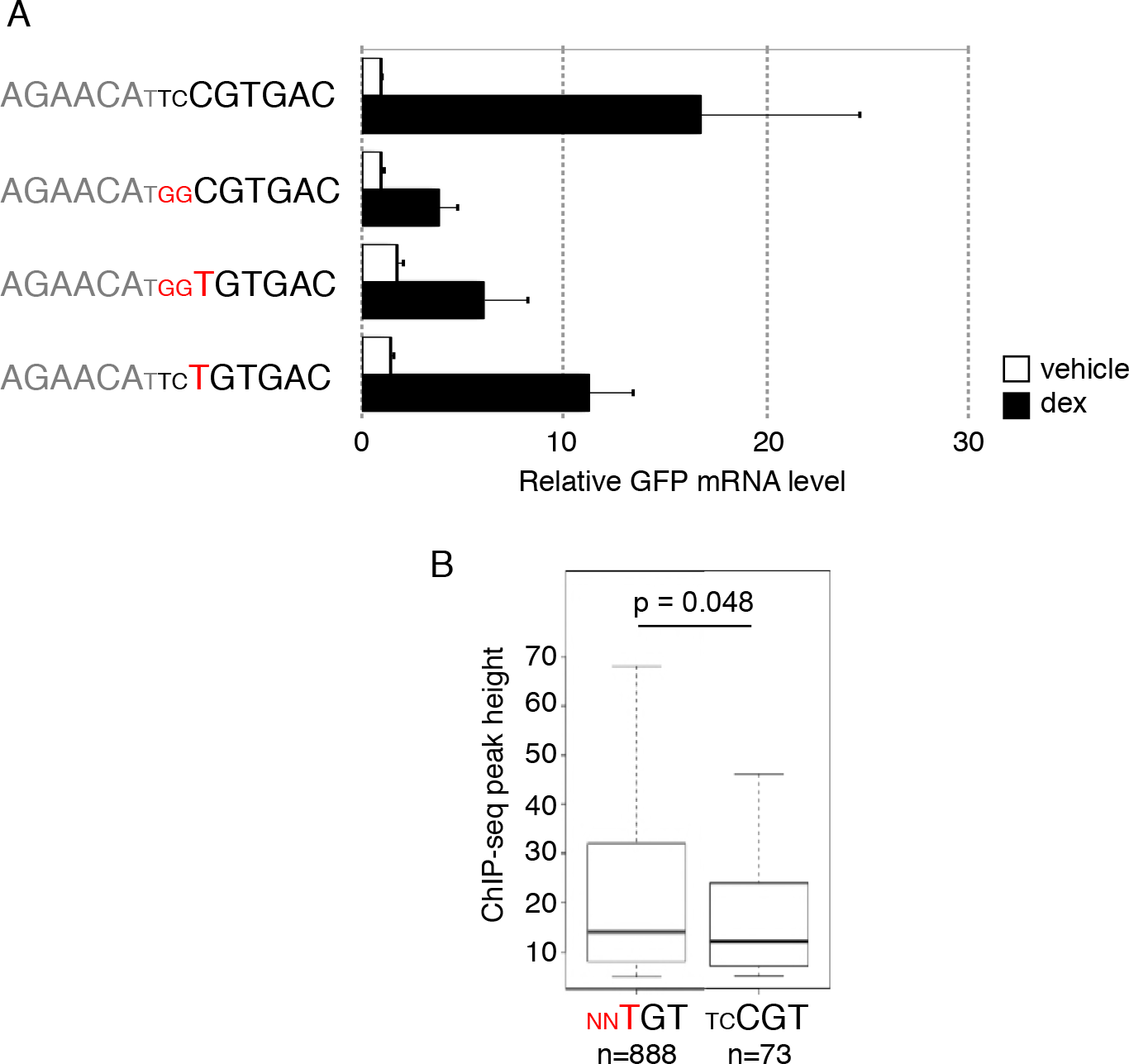
Characterization of the combi2 motif. (a) Transcriptional activity of STARR-seq reporters containing candidate GBS variants as indicated. Relative RNA levels ± S.E.M. are shown for cells treated with ethanol vehicle and for cells treated overnight with 1 μM dexamethasone (n = 3). (b) Boxplot showing the peak-height for GR target genes with either a canonical GBS motif match (nnTGT) or a combi2 motif match (tcCGT). Center lines show the median, p value was calculated using a Wilcoxon rank-sum test.

**Figure S9.**
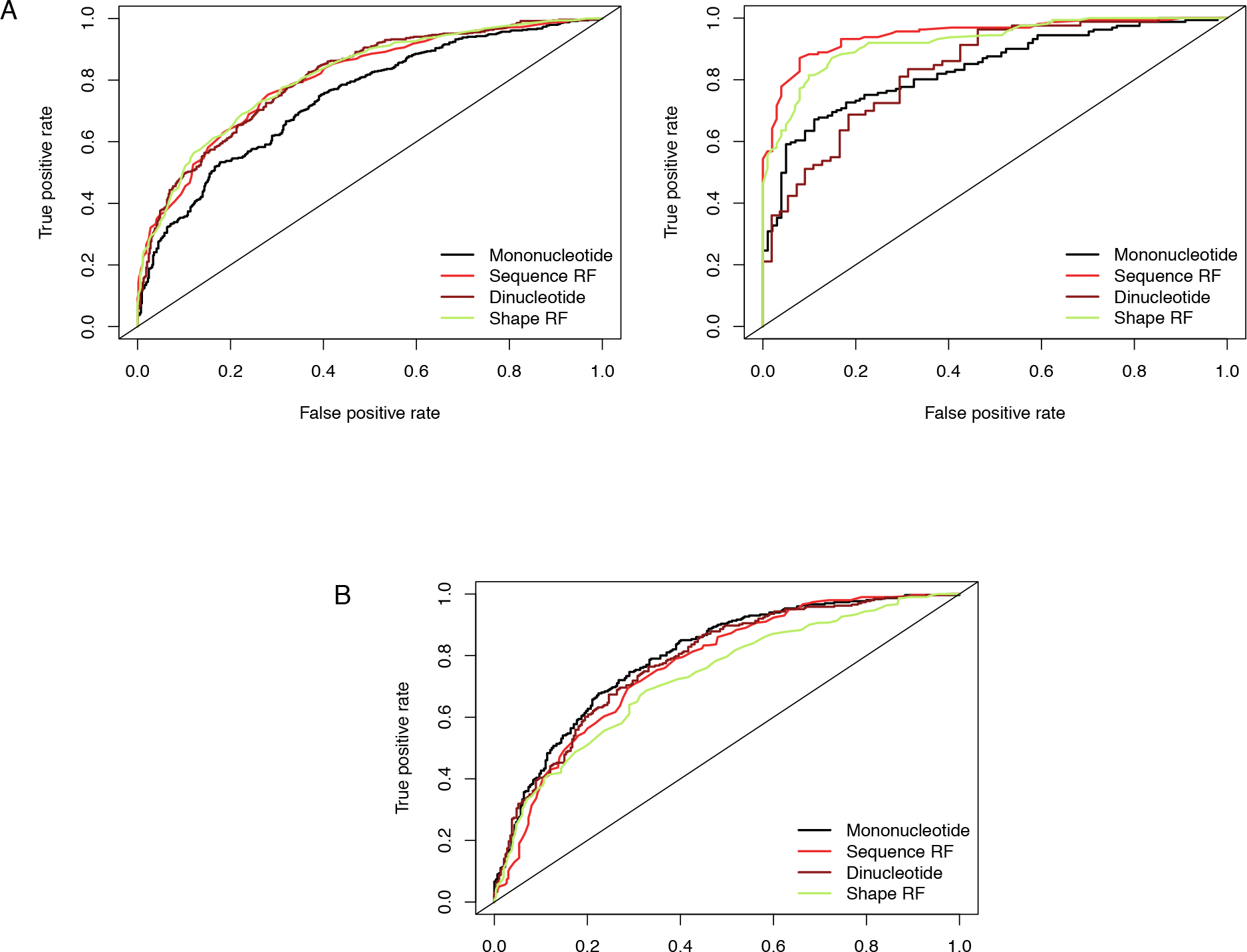
Prediction of GBS activity based on DNA sequence or DNA shape. (a) ROC curves analyzing the ability of the models to distinguish between blunting and enhancing flank variants for (left) the Cgt flank library; (right) the Sgk flank library. Mononucleotide: Classifier based on mononucleotide frequencies within the two classes. Dinucleotide: Classifier constructed using dinucleotide frequencies.Sequence Random Forest (RF): Random Forest classifer trained and tested on coded nucleotide sequences. Shape Random Forest (RF): Random forest classifier based on predicted MGW. (b) Same as for (a) except that model and ROC curves where trained and assessed for their ability to discriminate between the top and bottom 20% significantly active GBS variants from the half site library.Synthetic STARR-seq reveals how DNA shape and sequence modulate transcriptional output and noise.

## References

1. Grossman SR, Zhang X, Wang L, Engreitz J, Melnikov A, Rogov P, Tewhey R, Isakova A, Deplancke B, Bernstein BE, et al: Systematic dissection of genomic features determining transcription factor binding and enhancer function. Proc Natl Acad Sci U S A 2017, 114:E1291–E1300.

2. Schmid W, Strahle U, Schutz G, Schmitt J, Stunnenberg H: Glucocorticoid receptor binds cooperatively to adjacent recognition sites. EMBO J 1989, 8:2257–2263.

3. Schone S, Jurk M, Helabad MB, Dror I, Lebars I, Kieffer B, Imhof P, Rohs R, Vingron M, Thomas-Chollier M, Meijsing SH: Sequences flanking the core-binding site modulate glucocorticoid receptor structure and activity. Nat Commun 2016, 7:12621.

4. Watson LC, Kuchenbecker KM, Schiller BJ, Gross JD, Pufall MA, Yamamoto KR: The glucocorticoid receptor dimer interface allosterically transmits sequence-specific DNA signals. Nat Struct Mol Biol 2013, 20:876–883.

5. Meijsing SH, Pufall MA, So AY, Bates DL, Chen L, Yamamoto KR: DNA binding site sequence directs glucocorticoid receptor structure and activity. Science 2009, 324:407–410.

6. Maheshri N, O’shea EK: Living with noisy genes: how cells function reliably with inherent variability in gene expression. Annu Rev Biophys Biomol Struct 2007, 36:413–434.

7. Blake WJ, M Ka, Cantor CR, Collins JJ: Noise in eukaryotic gene expression. Nature 2003, 422:633–637.

8. Kaern M, Elston TC, Blake WJ, Collins JJ: Stochasticity in gene expression: from theories to phenotypes. Nat Rev Genet 2005, 6:451–464.

9. Raj A, van Oudenaarden A: Single-molecule approaches to stochastic gene expression. Annu Rev Biophys 2009, 38:255–270.

10. Ross IL, Browne CM, Hume DA: Transcription of individual genes in eukaryotic cells occurs randomly and infrequently. Immunol Cell Biol 1994, 72:177–185.

11. Suter DM, Molina N, Gatfield D, Schneider K, Schibler U, Naef F: Mammalian genes are transcribed with widely different bursting kinetics. Science 2011, 332:472–474.

12. Fritzsch C, Baumgartner S, Kuban M, Steinshorn D, Reid G, Legewie S: Estrogen-dependent control and cell-to-cell variability of transcriptional bursting. Mol Syst Biol 2018, 14:e7678.

13. Cai L, Friedman N, Xie XS: Stochastic protein expression in individual cells at the single molecule level. Nature 2006, 440:358–362.

14. Raser JM, O’Shea EK: Control of stochasticity in eukaryotic gene expression. Science 2004, 304:1811–1814.

15. Blake WJ, Balazsi G, Kohanski MA, Isaacs FJ, Murphy KF, Kuang Y, Cantor CR, Walt DR, Collins JJ: Phenotypic consequences of promoter-mediated transcriptional noise. Mol Cell 2006, 24:853–865.

16. Sharon E, van Dijk D, Kalma Y, Keren L, Manor O, Yakhini Z, Segal E: Probing the effect of promoters on noise in gene expression using thousands of designed sequences. Genome Res 2014, 24:1698–1706.

17. Hornung G, Bar-Ziv R, Rosin D, Tokuriki N, Tawfik DS, Oren M, Barkai N: Noise-mean relationship in mutated promoters. Genome Res 2012, 22:2409–2417.

18. Dey SS, Foley JE, Limsirichai P, Schaffer DV, Arkin AP: Orthogonal control of expression mean and variance by epigenetic features at different genomic loci. Mol Syst Biol 2015, 11:806.

19. Dadiani M, van Dijk D, Segal B, Field Y, Ben-Artzi G, Raveh-Sadka T, Levo M, Kaplow I, Weinberger A, Segal E: Two DNA-encoded strategies for increasing expression with opposing effects on promoter dynamics and transcriptional noise. Genome Res 2013, 23:966–976.

20. Raser JM, O’shea EK: Noise in gene expression: origins, consequences, and control. Science 2005, 309:2010–2013.

21. Inoue F, Ahituv N: Decoding enhancers using massively parallel reporter assays. Genomics 2015, 106:159–164.

22. Arnold CD, Gerlach D, Stelzer C, Boryn LM, Rath M, Stark A: Genome-wide quantitative enhancer activity maps identified by STARR-seq. Science 2013, 339:1074–1077.

23. Liu Y, Yu S, Dhiman VK, Brunetti T, Eckart H, White KP: Functional assessment of human enhancer activities using whole-genome STARR-sequencing. Genome Biol 2017, 18:219.

24. Vanhille L, Griffon A, Maqbool MA, Zacarias-Cabeza J, Dao LT, Fernandez N, Ballester B, Andrau JC, Spicuglia S: High-throughput and quantitative assessment of enhancer activity in mammals by CapStarr-seq. Nat Commun 2015, 6:6905.

25. Vockley CM, D’Ippolito AM, McDowell IC, Majoros WH, Safi A, Song L, Crawford GE, Reddy TE: Direct GR Binding Sites Potentiate Clusters of TF Binding across the Human Genome. Cell 2016, 166:1269–1281 e1219.

26. Shlyueva D, Stelzer C, Gerlach D, Yanez-Cuna JO, Rath M, Boryn LM, Arnold CD, Stark A: Hormone-responsive enhancer-activity maps reveal predictive motifs, indirect repression, and targeting of closed chromatin. Mol Cell 2014, 54:180–192.

27. Rogatsky I, Trowbridge JM, Garabedian MJ: Glucocorticoid receptor-mediated cell cycle arrest is achieved through distinct cell-specific transcriptional regulatory mechanisms. Mol Cell Biol 1997, 17:3181–3193.

28. Love MI, Huber W, Anders S: Moderated estimation of fold change and dispersion for RNA-seq data with DESeq2. Genome Biol 2014, 15:550.

29. Wu X, Bartel DP: kpLogo: positional k-mer analysis reveals hidden specificity in biological sequences. Nucleic Acids Res 2017, 45:W534–W538.

30. Zhang L, Martini GD, Rube HT, Kribelbauer JF, Rastogi C, FitzPatrick VD, Houtman JC, Bussemaker HJ, Pufall MA: SelexGLM differentiates androgen and glucocorticoid receptor DNA-binding preference over an extended binding site. Genome Res 2018, 28:111–121.

31. Chiu TP, Comoglio F, Zhou T, Yang L, Paro R, Rohs R: DNAshapeR: an R/Bioconductor package for DNA shape prediction and feature encoding. Bioinformatics 2016, 32:1211–1213.

32. Starick SR, Ibn-Salem J, Jurk M, Hernandez C, Love MI, Chung HR, Vingron M, Thomas-Chollier M, Meijsing SH: ChIP-exo signal associated with DNA-binding motifs provides insight into the genomic binding of the glucocorticoid receptor and cooperating transcription factors. Genome Res 2015, 25:825–835.

33. Rhee HS, Pugh BF: Comprehensive genome-wide protein-DNA interactions detected at singlenucleotide resolution. Cell 2011, 147:1408–1419.

34. Strahle U, Schmid W, Schutz G: Synergistic action of the glucocorticoid receptor with transcription factors. EMBO J 1988, 7:3389–3395.

35. Pearce D, Yamamoto KR: Mineralocorticoid and glucocorticoid receptor activities distinguished by nonreceptor factors at a composite response element. Science 1993, 259:1161–1165.

36. Yang L, Orenstein Y, Jolma A, Yin Y, Taipale J, Shamir R, Rohs R: Transcription factor family-specific DNA shape readout revealed by quantitative specificity models. Mol Syst Biol 2017, 13:910.

37. Rohs R, Jin X, West SM, Joshi R, Honig B, Mann RS: Origins of specificity in protein-DNA recognition. Annu Rev Biochem 2010, 79:233–269.

38. Abe N, Dror I, Yang L, Slattery M, Zhou T, Bussemaker HJ, Rohs R, Mann RS: Deconvolving the recognition of DNA shape from sequence. Cell 2015, 161:307–318.

39. Zheng J, Chang MR, Stites RE, Wang Y, Bruning JB, Pascal BD, Novick SJ, Garcia-Ordonez RD, Stayrook KR, Chalmers MJ, et al: HDX reveals the conformational dynamics of DNA sequence specific VDR co-activator interactions. Nat Commun 2017, 8:923.

40. Zhang J, Chalmers MJ, Stayrook KR, Burris LL, Wang Y, Busby SA, Pascal BD, Garcia-Ordonez RD, Bruning JB, Istrate MA, et al: DNA binding alters coactivator interaction surfaces of the intact VDR-RXR complex. Nat Struct Mol Biol 2011, 18:556–563.

41. Hall JM, McDonnell DP, Korach KS: Allosteric regulation of estrogen receptor structure, function, and coactivator recruitment by different estrogen response elements. Mol Endocrinol 2002, 16:469–486.

42. Parker SC, Hansen L, Abaan HO, Tullius TD, Margulies EH: Local DNA topography correlates with functional noncoding regions of the human genome. Science 2009, 324:389–392.

43. Mathelier A, Xin B, Chiu TP, Yang L, Rohs R, Wasserman WW: DNA Shape Features Improve Transcription Factor Binding Site Predictions In Vivo. Cell Syst 2016, 3:278–286 e274.

44. Singh A, Razooky B, Cox CD, Simpson ML, Weinberger LS: Transcriptional bursting from the HIV-1 promoter is a significant source of stochastic noise in HIV-1 gene expression. Biophys J 2010, 98:L32–34.

45. Dobin A, Davis CA, Schlesinger F, Drenkow J, Zaleski C, Jha S, Batut P, Chaisson M, Gingeras TR: STAR: ultrafast universal RNA-seq aligner. Bioinformatics 2013, 29:15–21.

46. Crooks GE, Hon G, Chandonia JM, Brenner SE: WebLogo: a sequence logo generator. Genome Res 2004, 14:1188–1190.

47. Thomas-Chollier M, Defrance M, Medina-Rivera A, Sand O, Herrmann C, Thieffry D, van Helden J: RSAT 2011: regulatory sequence analysis tools. Nucleic Acids Res 2011, 39:W86–91.

48. Turatsinze JV, Thomas-Chollier M, Defrance M, van Helden J: Using RSAT to scan genome sequences for transcription factor binding sites and cis-regulatory modules. Nat Protoc 2008, 3:1578–1588.

49. Wiener ALaM: Classification and Regression by randomForest. R News 2002, 2:18–22.

50. Khan A, Fornes O, Stigliani A, Gheorghe M, Castro-Mondragon JA, van der Lee R, Bessy A, Cheneby J, Kulkarni SR, Tan G, et al: JASPAR 2018: update of the open-access database of transcription factor binding profiles and its web framework. Nucleic Acids Res 2018, 46:D260–D266.

51. van Dijk M, Bonvin AM: 3D-DART: a DNA structure modelling server. Nucleic Acids Res 2009, 37:W235–239.

52. Jia Y, Dewey TG, Shindyalov IN, Bourne PE: A new scoring function and associated statistical significance for structure alignment by CE. J Comput Biol 2004, 11:787–799.

